# A toolbox of astrocyte-specific, serotype-independent adeno-associated viral vectors using microRNA targeting sequences

**DOI:** 10.1101/2023.02.21.529451

**Authors:** Amy J. Gleichman, Riki Kawaguchi, Michael V. Sofroniew, S. Thomas Carmichael

## Abstract

Astrocytes, one of the most prevalent cell types in the central nervous system (CNS), are critically involved in neural function in both health and disease. Genetically manipulating astrocytes is an essential tool in understanding and affecting their roles. Adeno-associated viruses (AAVs) enable rapid genetic manipulation; however, astrocyte specificity of AAVs can be limited, with high off-target expression in neurons and sparsely in endothelial cells. Here, we report the development of a cassette of four copies of six miRNA targeting sequences (4×6T) which triggers transgene degradation specifically in neurons and endothelial cells. When used in combination with the GfaABC1D promoter, 4×6T increases astrocytic specificity of Cre with a viral reporter from <50% to >99% in multiple serotypes in mice, and confers astrocyte specificity in two inducible forms of Cre; Dre; and reporters. We also present empty vectors to add 4×6T to other cargo, independently and in Cre/Dre-dependent forms. This toolbox of AAVs provides a way to rapidly manipulate astrocytes throughout the CNS, is compatible with different AAV serotypes, and demonstrates the efficacy of using multiplexed miRNA targeting sequences to decrease expression in multiple off-target cell populations simultaneously.

---

Astrocytes comprise roughly 20-40% of the cells in the CNS and are found across all brain regions^1^. They play crucial roles in both physiological and pathological states and across the lifespan^2^. The ability to specifically genetically manipulate astrocytes is critical in order to understand and alter astrocytic functions; to this end, viral vectors can offer experimental flexibility, speed, and selectivity^3^. Adeno-associated viruses (AAVs) are widely used viral vectors due to their safety profile, limited toxicity, and the natural occurrence of serotypes with different tropism. Furthermore, engineered AAV capsids can confer new properties on the virus. For example, newly developed AAV capsids such as PHP.eB^4^ are systemically deliverable; these viruses can be injected into mice intravenously (IV), cross the blood-brain barrier, and transduce cells throughout the CNS with a single injection.

Implementing these new vectors in astrocytes, however, is complicated by non-astrocytic expression, despite the use of astrocyte-specific promoters. This expression is variable and depends on the capsid used^5^ and the cargo expressed^6,7^, but extensive neuronal expression can be observed and complicates the interpretation of results^6,8–10^. This is particularly problematic when delivering Cre recombinase, which requires only low expression levels for effective genetic recombination. One way to obtain serotype-independent specificity in viral vectors - that is, specificity that is compatible with different capsids - is through the use of microRNA (miRNA) targeting sequences^11^.

These targeting sequences are DNA sequences that will bind to specific miRNAs that are highly expressed in off-target cells but not in the cells of interest: in off-target cells, transgene mRNA will bind miRNAs and be degraded, but will be preserved in target cells and allow expression. One benefit to this approach to cell-type specificity is that these sequences can be used within the recombinant AAV genome, rather than the capsid, and therefore can improve specificity across serotypes. Using miR-124, this approach increased astrocyte selectivity in lentiviruses^12^ and in AAVs for expression of GFP^8,13^ and Cre^8^. While miR-124 targeting improves astrocyte specificity, off-target expression in neurons and endothelial cells was still readily observed, particularly with systemic Cre delivery^8^. This suggests that while miR targeting holds promise for astrocytic viral specificity, there is room for improvement, particularly when delivering recombinases.

Here, we developed a cassette of miRNA targeting sequences, comprised of four copies of each of six miRNAs, to de-target neurons and endothelial cells simultaneously and improve astrocytic specificity in AAV vectors across multiple serotypes and a variety of cargo, including multiple recombinases. This combined toolbox of vectors presents improved options for astrocyte-specific viral manipulation.

## Results

### Development of miR targeting cassette to improve astrocyte specificity

To assess the astrocytic specificity of Cre viral delivery using current vectors for systemic delivery, we administered PHP.eB viruses carrying Cre under a glial fibrillary acidic protein (GFAP) promoter (PHP.eB::GFAP-Cre; Addgene 105550); GFAP is an astrocyte-specific protein and elements of the promoter have been widely used to target astrocytes^14^. We also generated a vector using the smaller GfaABC1D promoter^15^ (PHP.eB::GfaABC1D-Cre). As only low levels of Cre are necessary for recombination, we omitted a woodchuck hepatitis virus post-transcriptional regulatory element (WPRE), used to increase transgene expression, reasoning that this might decrease off-target transgene expression; a similar approach was recently used to improve targeting of Muller glia^16^. Viruses were delivered systemically using retroorbital administration into young adult (2-5 month) C57BL/6J mice. To assess Cre activity, we co-injected a Cre-dependent GFP reporter (PHP.eB::CAG-flex-GFP, Addgene 28304); two weeks post-injection, we evaluated colocalization of GFP^+^ cells, enhanced with an anti-GFP antibody, with the astrocytic nuclear marker Sox9^17^ in the cortex. Both promoters were predominantly non-astrocytic, although GfaABC1D showed higher astrocyte selectivity (Fig. 1a; Sox9^+^ GFAP: 22.79% ± 1.50 vs GfaABC1D: 38.81% ± 3.64); the majority of Sox9^-^GFP^+^ cells were neurons (NeuN^+^). Viral Cre-dependent flex constructs can show Cre-independent expression^18^; therefore, we repeated PHP.eB::GfaABC1D-Cre delivery in Ai14-tdTomato mice^19^, a transgenic line with Cre-dependent tdTomato expression. While astrocyte specificity was greater in these mice, we still observed extensive neuronal (NeuN^+^tdTomato^+^, 29.16% ± 2.78) and minimal endothelial (CD31^+^ tdTomato^+^, 1.23% ± 0.07) contamination (Fig. 1b).

**Figure 1.**
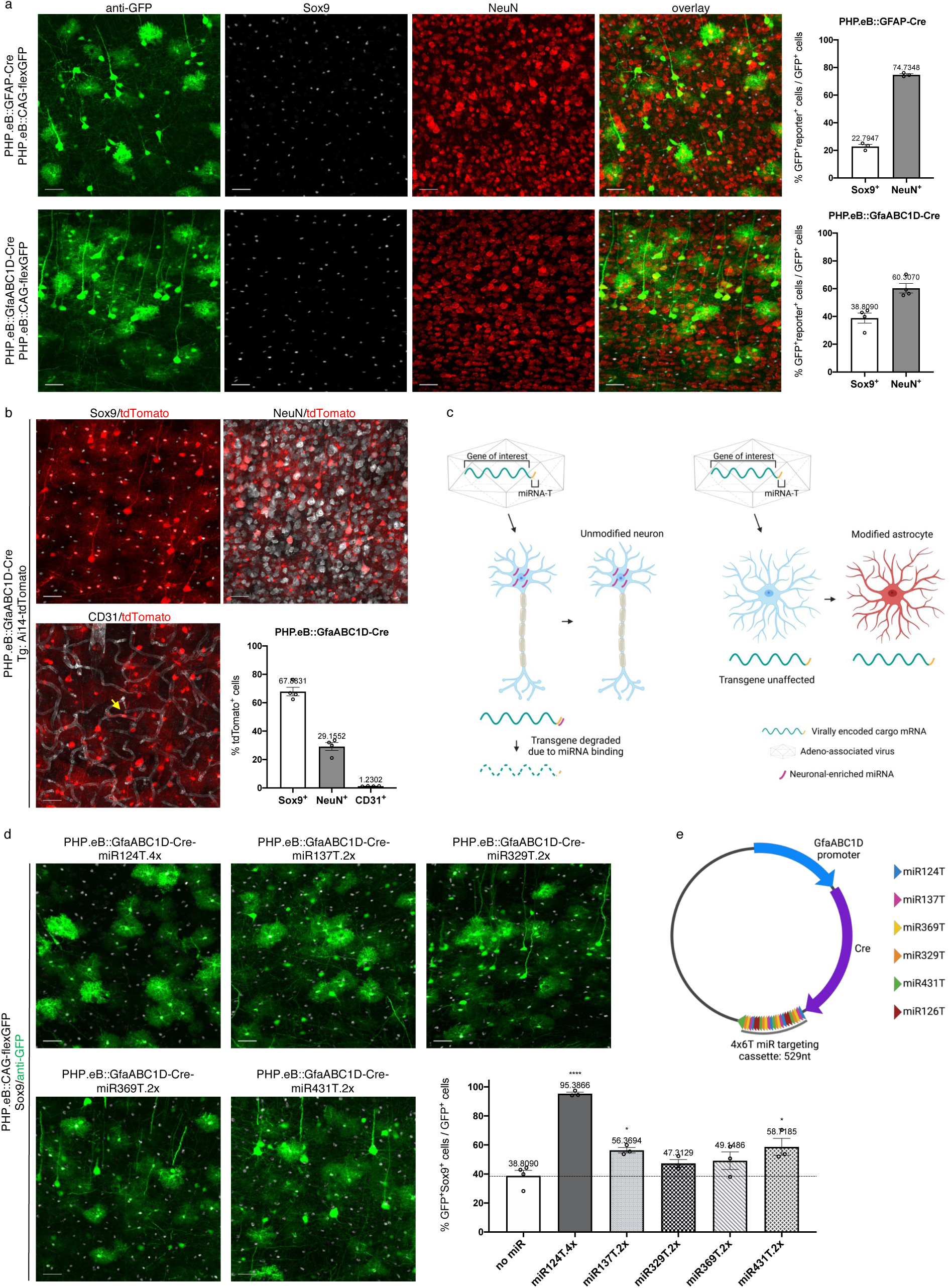
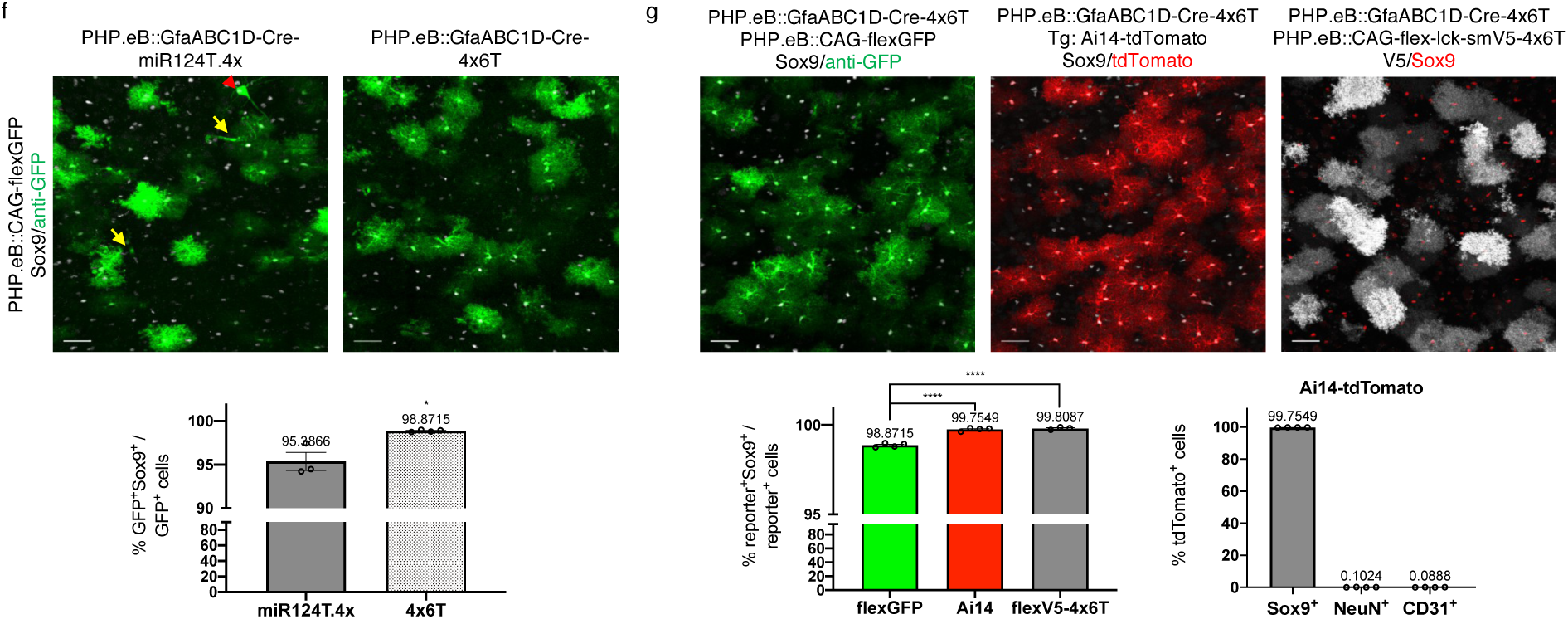
Low astrocytic viral specificity can be enhanced with miRNA targeting sequences. **a**, Systemic delivery of two different GFAP-based promoters (GFAP, n=3 mice; and GfaABC1D, n=4 mice), co-injected with PHP.eB::CAG-flex-GFP, yield high levels of transduction of non-astrocytic cells. Sox9, astrocytes; NeuN, neurons. **b**, Characterization of PHP.eB::GfaABC1D-Cre delivery in transgenic (Tg) Ai14-tdTomato mice: while the majority of tdTomato^+^ cells are astrocytes, there are high numbers of transduced neurons (NeuN^+^) and low numbers of endothelial cells (CD31^+^; yellow arrow); n=4 mice. **c**, Schematic diagram of miRNA targeting approach. **d**, Changes in astrocytic selectivity by the addition of multiple copies of single miR targeting sequences (n=3 mice); compared to GfaABC1D-Cre (n=4 mice) (one-way ANOVA, Dunnett’s multiple comparisons test, p<0.0001, F=24.76, df=18. All vs no miR: miR124T.4×, four copies, ****p<0.0001; miR137T.2×, two copies, *p=0.0263; miR329T.2×, two copies, p=0.4348; miR369T.2×, two copies, p=0.2680; miR431T.2×, two copies, *p=0.0118.) Dotted line denotes astrocyte specificity in GfaABC1D-Cre with no miR cassettes. **e**, Schematic diagram of final AAV::GfaABC1D-Cre-4×6T plasmid. **f**, Enhanced astrocyte specificity with PHP.eB::GfaABC1D-Cre-4×6T (n=4 mice) vs PHP.eB::GfaABC1D-Cre-miR124T.4× (n=3 mice), using a PHP.eB::CAG-flex-GFP reporter (two-tailed unpaired t-test, *p=0.0103, t=4.002, df=5). Non-astrocytes expressing GFP after transduction with PHP.eB::GfaABC1D-Cre-miR124T.4× include both neurons (red arrowhead) and endothelial cells (yellow arrows). **g**, Further enhancement of astrocyte specificity using PHP.eB::GfaABC1D-Cre-4×6T with a transgenic Ai14-tdTomato reporter (n=4 mice) or a PHP.eB::CAG-flex-smV5-4×6T reporter (n=3 mice) rather than PHP.eB::CAG-flex-GFP (n=4 mice). Using a transgenic mouse line or including the 4×6T cassette on the flex reporter increases astrocyte specificity (one-way ANOVA, Tukey’s multiple comparisons test, p=0.0144, F=7.548, df=10; flexGFP vs Ai14, *p=0.0249; flexGFP vs flexV5-4×6T, *p=0.0259). All data presented as mean ± SEM. Scale bars: 40μm.

To decrease neuronal expression, we added a previously reported cassette with four copies of miR-124-3p targeting sequence (miR124T.4×^12^) immediately following the stop codon terminating GfaABC1D-Cre (Fig. 1c). We also identified four other miRNAs –miR-137-3p, miR-329-3p, miR-369-5p, and miR-431-3p – with similar patterns of high neuronal/low astrocytic expression^20,21^ and generated viral vectors for each that included two copies of the targeting sequence. We delivered these vectors systemically using PHP.eB, co-injecting with PHP.eB::CAG-flex-GFP, and co-stained for GFP and Sox9. As expected, miR124T.4× showed high levels of astrocyte specificity (95.39% ± 1.038 Sox9^+^). miR137T.2× and miR431T.2× showed lower, but significant, increases in astrocyte specificity (miR137T.2×: 56.37% ± 1.85 Sox9^+^; miR431T.2×: 58.72% ± 5.85 Sox9^+^). While miR329T.2× and miR369T.2× did not show a significantly different proportion of astrocytic specificity, they appeared to trend towards greater specificity (miR329T.2×: 47.31% ± 2.62 Sox9^+^; miR369T.2×: 49.15% ± 6.06 Sox9^+^) (Fig. 1d); notably, these increases in specificity seen with the four previously untested miRs were observed with only two copies of their relative targeting sequences, to facilitate rapid sequence synthesis. Therefore, in order to target astrocytes as specifically and robustly as possible, we developed a multiplexed vector with four interspersed copies of the targeting sequences for each miR. To de-target endothelial cells, we also included four copies of a targeting sequence for miR-126a-3p^22^, generating a final cassette of four copies of each of six miRNA targeting sequences with a final length of 529 nucleotides (4×6T, Fig. 1e).

We systemically injected PHP.eB::GfaABC1D-Cre-4×6T into wildtype mice with PHP.eB::CAG-flex-GFP (5×10^11vg/mouse per virus) and evaluated astrocyte specificity in the cortex. Compared to PHP.eB::GfaABC1D-Cre-miR124T.4×, the presence of additional miRNA targeting cassettes significantly improved astrocyte specificity (Fig. 1f; miR124T.4×, 95.39% ± 1.04 Sox9^+^ vs 4×6T, 98.87% ± 0.06 Sox9^+^). To test the specificity of PHP.eB::GfaABC1D-Cre-4×6T in transgenic animals, we injected virus into Ai14-tdTomato mice (5×10^11vg/mouse); this yielded even higher astrocyte specificity compared to PHP.eB::CAG-flex-GFP reporter (Fig. 1g; Ai14: 99.75% ± 0.04 Sox9^+^ vs flex-GFP, 98.87% ± 0.06 Sox9^+^). This reflected successful de-targeting of both neurons and endothelial cells (Fig. 1g; 284.7-fold reduction in % tdTomato^+^NeuN^+^/tdTomato^+^ cells; 13.9-fold reduction in % tdTomato^+^CD31^+^/tdTomato^+^ cells). Further, the relatively lower astrocyte specificity seen with PHP.eB::CAG-flex-GFP compared to Ai14-tdTomato could be improved by adding the 4×6T cassette to a viral flex reporter (membrane-targeted V5 spaghetti monster reporter^23^, PHP.eB::CAG-flex-lck-smV5-4×6T; “flexV5-4×6T”) (Fig. 1g; 99.81% ± 0.04 Sox9 ^+^ cells), thereby adding miR targeting specificity to both the Cre and reporter components of the expression system.

### Role of titer and serotype in maximizing astrocyte specificity

While astrocyte labeling with PHP.eB::GfaABC1D-Cre-4×6T was highly specific, it did not transduce all astrocytes. To improve the percentage of astrocytes labeled, we injected PHP.eB::GfaABC1D-Cre-4×6T systemically at a higher titer (3×10^12 vg/mouse). This titer maintained 99.67% ± 0.12 astrocyte specificity while labeling 65.66% ± 1.38 of astrocytes in the cortex (Fig. 2a). To label a greater percentage in a specific area, we injected virus intracortically (Fig. 2b). However, intracortical injection of high titer (500nl of 1×10^12 vg/ml per virus) of PHP.eB::GfaABC1D-Cre-4×6T with PHP.eB::CAG-flex-lck-smV5-4×6T showed extensive neuronal transduction (Fig. 2c), despite the presence of the 4×6T cassette on both viruses. Therefore, we repackaged the reporter construct in the more astrocyte-selective AAV2/5 serotype^5^ (AAV2/5::CAG-flex-lck-smV5-4×6T) and co-injected PHP.eB::GfaABC1D-Cre-4×6T and AAV2/5::CAG-flex-lck-smV5-4×6T into Ai14-tdTomato mice. High-titer intracortical injection transduced all astrocytes within the core of the injected region, but with relatively lower astrocyte specificity (Fig 2d, 87.19% ± 3.97 tdTomato^+^Sox9^+^/tdTomato^+^). Using the AAV2/5 reporter, however, maintained astrocyte selectively even at high titers (Fig. 2d, 99.75% ± 0.09 V5^+^Sox9^+^/V5^+^). Thus, it is possible to label a high percentage of geographically restricted astrocytes through direct injection of virus into the brain by combining capsid choice and the inclusion of the miR targeting cassette on the delivered gene.

**Figure 2.**
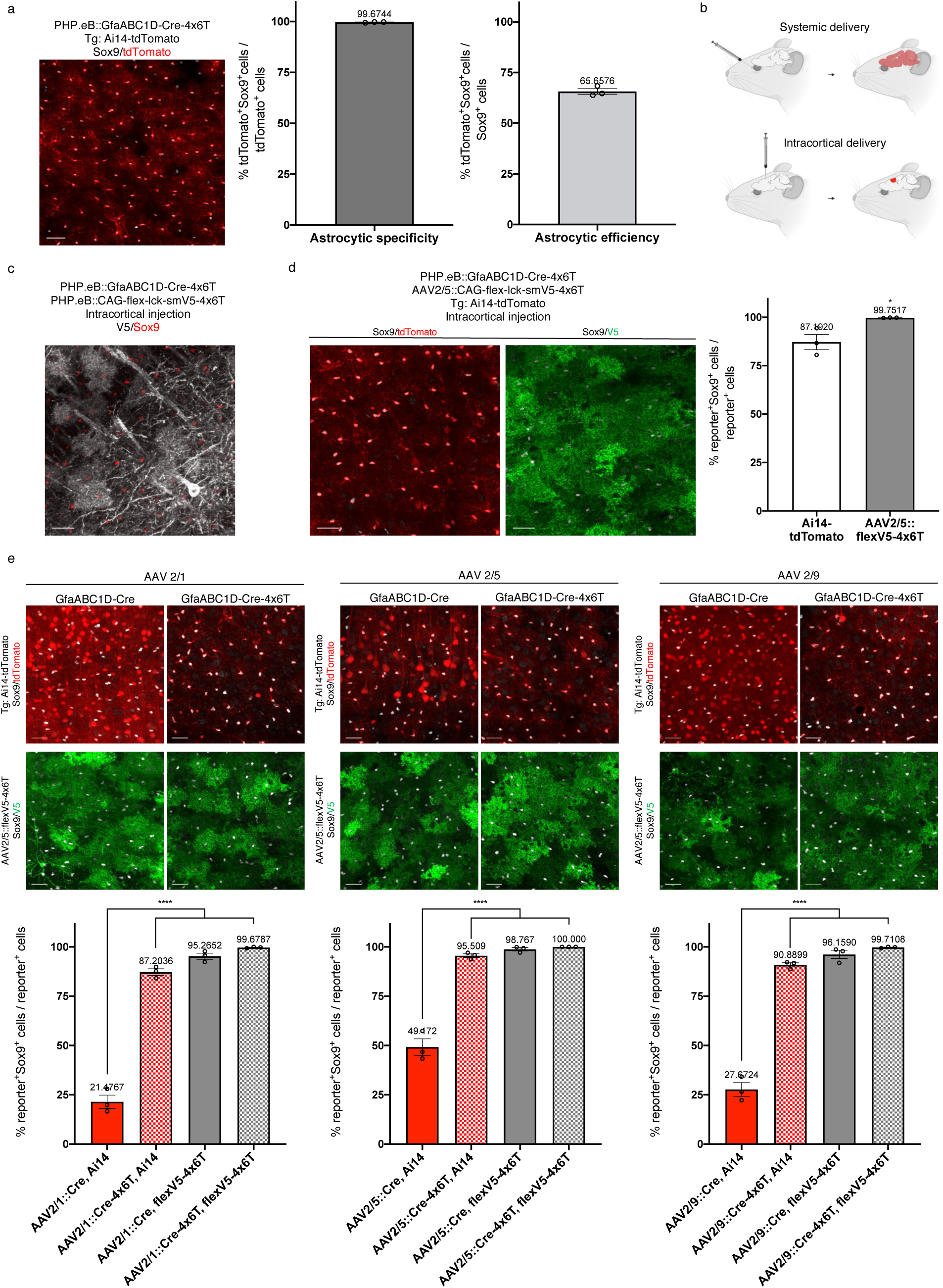
4×6T cassette increases astrocyte specificity at high titers and across serotypes. **a**, High-titer (3×10^12 vg/mouse) systemic delivery of PHP.eB::GfaABC1D-Cre-4×6T in Ai14-tdTomato mice yields 99.67% astrocyte specificity (% of tdTomato^+^ cells that were Sox9^+^); the efficiency of transduction (% of Sox9^+^ cells transduced) was 65.66% of Sox9^+^ cells in the cortex. Scale bar: 40μm. **b**, Schematic of viral delivery approaches: retroorbital injection leads to lower levels of brain-wide transduction, while direct intracortical injection leads to higher levels of transduction within a narrow region around the injection site. **c**, High-titer direct intracortical injection (500nl of 1×10^12 vg/ml per virus) of PHP.eB::GfaABC1D-Cre-4×6T and PHP.eB::CAG-flex-lck-smV5-4×6T shows high levels of neuronal contamination. Scale bar: 30μm. **d**, High-titer direct intracortical injection of PHP.eB::GfaABC1D-Cre-4×6T and AAV2/5::CAG-flex-lck-smV5-4×6T into Ai14-tdTomato mice shows predominantly astrocytic expression of the tdTomato reporter but more astrocyte-specific expression of the AAV2/5::flexV5-4×6T reporter. Two-tailed unpaired t-test, *p=0.0340; t=3.165, df=4. Scale bars: 30μm. **e**, High-titer direction intracortical injection of GfaABC1D-Cre-4×6T packaged in different serotypes, co-injected with AAV2/5::CAG-flex-lck-smV5-4×6T into Ai14-tdTomato mice. Serotype impacts astrocyte specificity, with highest specificity with AAV2/5. The presence of the 4×6T on Cre, on the reporter virus, or both increases astrocytic specificity, with the highest specificity seen when both elements of a Cre/reporter system are tagged with 4×6T. Within serotypes: GfaABC1D-Cre vs all 4×6T conditions, one-way ANOVA with Tukey’s multiple comparisons test, ****p<0.0001. AAV2/1: F=320.2, df=11. AAV2/5: F=124.4, df=11. AAV2/9: F=259.0, df=11. Scale bars: 30μm. All data presented as mean ± SEM; n=3 mice per condition.

To evaluate more broadly how serotype affects astrocyte specificity with 4×6T, we packaged GfaABC1D-Cre and GfaABC1D-Cre-4×6T in AAV2/1, AAV2/5, and AAV2/9, and injected each virus intracortically with AAV2/5::CAG-flex-lck-smV5-4×6T into Ai14-tdTomato mice. In the absence of the 4×6T cassette, all Cre viruses showed high levels of tdTomato^+^ neuronal contamination, although AAV2/5 showed the highest astrocyte specificity (AAV2/1: 21.48% ± 3.39 Sox9^+^; AAV2/5: 49.17% ± 4.21 Sox9^+^; AAV2/9: 27.67% ± 3.49 Sox9^+^; Fig. 2e). Adding the 4×6T cassette to Cre increased astrocyte specificity of tdTomato expression across all serotypes (AAV2/1: 87.20% ± 1.67 Sox9^+^; AAV2/5: 95.51% ± 0.95 Sox9^+^; AAV2/9: 90.89% ± 1.11 Sox9^+^; Fig. 2e).

Including the 4×6T cassette on the viral reporter (AAV2/5::CAG-flex-lck-smV5-4×6T) improved astrocyte specificity when using GfaABC1D-Cre with no miR cassette (AAV2/1-Cre: 95.27% ± 1.57 Sox9^+^; AAV2/5-Cre: 98.77% ± 0.84 Sox9^+^; AAV2/9-Cre: 96.16% ± 2.13 Sox9^+^; Fig. 2e). This was further improved using both Cre-4×6T and the 4×6T reporter (AAV2/1-Cre-4×6T: 99.68% ± 0.19 Sox9^+^; AAV2/5-Cre-4×6T: 100% ± 0 Sox9^+^; AAV2/9-Cre-4×6T: 99.71% ± 0.21 Sox9^+^; Fig. 2e). AAV2/5 showed the highest level of astrocytic specificity in all conditions, suggesting that serotype can influence specificity, although the addition of the 4×6T cassette on both components of a fully viral Cre/flex system yields extremely high astrocyte specificity at high titers across multiple serotypes.

### 4×6T cassette confers astrocyte specificity across the lifespan and in reactive astrocytes

One advantage of using viral systems is flexibility in delivering virus at different ages. Therefore, we tested the astrocyte specificity of virus delivered across the lifespan of the mouse, administering virus systemically in pups at postnatal day 1-2 via temporal vein injection and in 28-month-old mice via retroorbital injection. We found that high astrocyte specificity conferred by the 4×6T cassette is maintained in the cortex across the mouse lifespan (P1-2: 99.68% ± 0.17 Sox9^+^; 28m: 99.76% ± 0.09 Sox9^+^; Fig 3a). However, one concern when using a GFAP-based promoter is that GFAP is also expressed in neural progenitor cells (NPCs); therefore, we evaluated astrocyte-specific expression in the dentate gyrus of the hippocampus, a region of ongoing neurogenesis in the mouse brain. Two weeks after virus delivery, we routinely found Sox9^+^V5^+^ radial glia in the hippocampus of P1-2 mice but not in young adult (2-5 month old) animals (Fig 3b).

**Figure 3.**
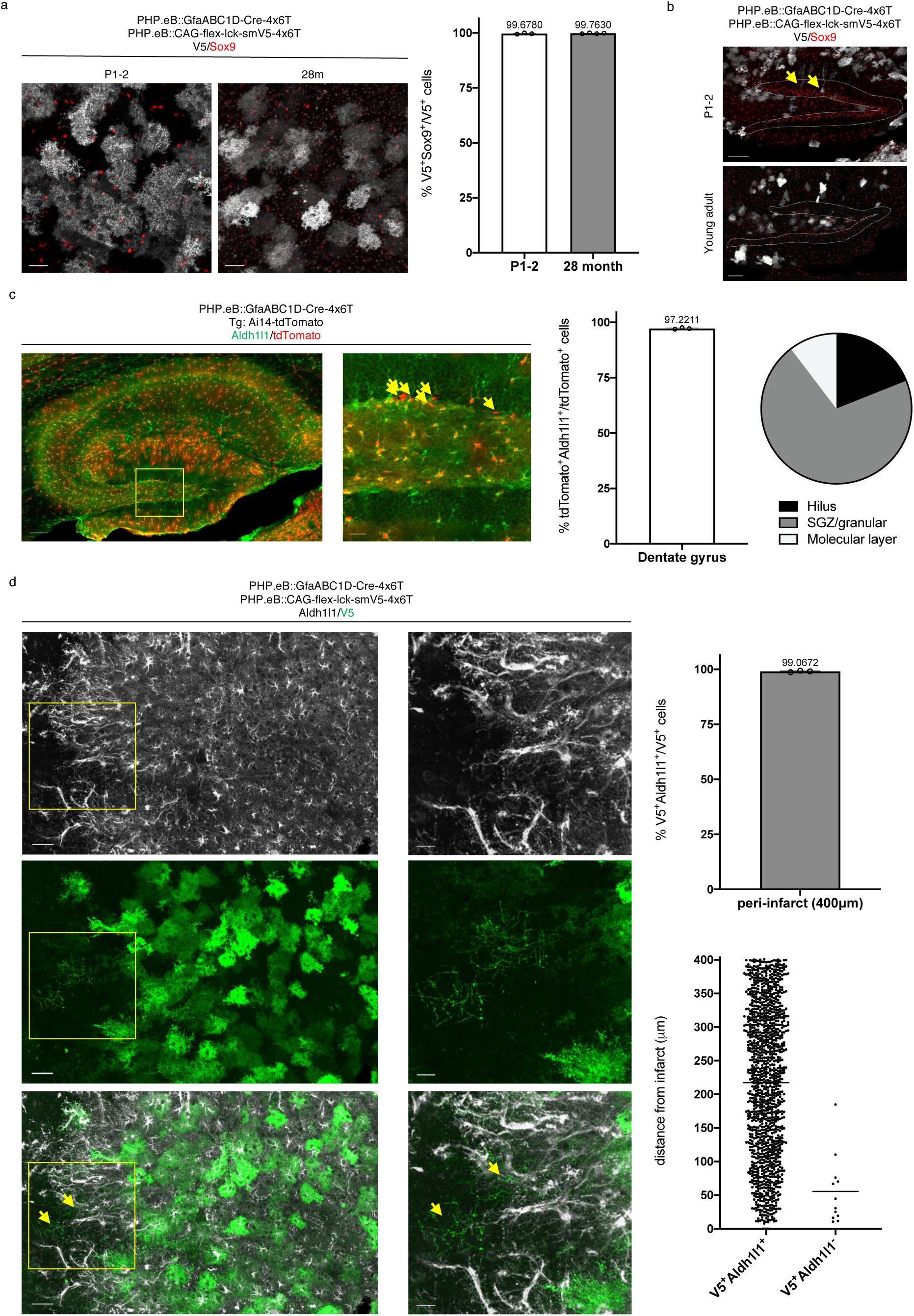
4×6T cassette confers astrocyte specificity across the lifespan and in reactive astrocytes. **a**, Systemic injection in mice as young as postnatal day 1-2 (temporal vein injection; 1×10^10 vg/mouse PHP.eB::GfaABC1D-Cre-4×6T + 2×10^10 vg/mouse PHP.eB::CAG-flex-lck-smV5-4×6T; n=3 mice) and as old as 28 months (retroorbital injection; 2×10^11 vg/mouse PHP.eB::GfaABC1D-Cre-4×6T and PHP.eB::CAG-flex-lck-smV5-4×6T; n=4 mice) show highly astrocyte-specific viral expression patterns. Scale bars: 40μm. **b**, Systemic injection in P1-2 pups yields transduced radial glia (yellow arrows) in the dentate gyrus, which is not readily observed in animals injected at 2-5 months; dotted gray lines, granule cell layer. Scale bars: 100μm. **c**, High-titer PHP.eB::GfaABC1D-Cre-4×6T injection (3×10^12 vg/mouse) in adult Ai14-tdTomato mice (n=3 mice) results in high astrocyte specificity in the dentate gyrus. The majority of Aldh1l1^-^tdTomato^+^ cells are found in the SGZ and granular layers (combined totals, 3 mice, 2 sections/mouse; total = 58 cells). Scale bars: left, 100μm; right, 20μm. **d**, 4×6T cassette maintains high levels of astrocytic specificity after stroke (n=3 mice), although morphologically distinct V5^+^Aldh1l1^-^ can be found near the infarct border. Left, 50μm scale bar; right, 20μm scale bar. Yellow box denotes area of higher magnification at right; yellow arrows indicate V5^+^Aldh1l1^-^ cells. V5^+^Aldh1l1^-^ cells: mean distance of 55 μm ± 14.81 from infarct border. Bar graphs presented as mean ± SEM.

To more fully evaluate astrocytic specificity in the dentate gyrus, we returned to high-titer injections (3×10^12 vg/mouse) of PHP.eB::GfaABC1D-Cre-4×6T in Ai14-tdTomato reporter mice. Because Sox9 is a marker of both astrocytes and progenitor populations, we instead used the cytosolic astrocytic marker Aldh1l1 to label astrocytes. While Aldh1l1 can also be found in neural stem cells^24^, Aldh1l1 expression in radial glia is relatively lower than Sox9^25^, and Aldh1l1-CreERT2 transgenic mice show minimal neural stem cell recombination in the dentate gyrus when tamoxifen is administered in adulthood^26^, suggesting it is a more accurate marker of astrocytic identity in neurogenic regions. We found a high degree of astrocyte specificity within the dentate gyrus (97.22% ± 0.21 Aldh1l1^+^; Fig 3c); those tdTomato^+^ cells that were Aldh1l1^-^ were found predominantly in the subgranular zone and granular layer (Fig 3c), as would be expected for radial glia and their progeny. The relatively sparse labeling of Aldh1l1^-^ cells in this neurogenic zone in adult mice suggests that the 4×6T cassette may de-target progenitor cells to some extent.

In response to CNS injury, astrocytes upregulate GFAP^27^ and can locally proliferate^28^. Injury can also induce proliferation and maturation of NPCs; in stroke, this results in new neurons in the peri-infarct cortex^29^. Therefore, we evaluated whether stroke affects astrocyte specificity of the 4×6T cassette. Young adult mice were systemically injected with PHP.eB::GfaABC1D-Cre-4×6T and PHP.eB::CAG-flex-lck-smV5-4×6T, received distal middle cerebral artery occlusion strokes (dMCAO) two weeks later, and were euthanized one week post-stroke; dMCAO produces a cortical infarct with extensive neurogenesis at this timepoint^29^. We examined astrocyte specificity of virally labeled cells in the cortex within 400μm of the infarct border, which encompasses essentially all proliferating astrocytes post-stroke^28^. Within the periinfarct cortex, we found that astrocyte specificity of viral expression was maintained to a high degree (99.07% ± 0.19 Aldh1l1^+^; Fig 3d); although we did find rare non-astrocytic, morphologically distinct Aldh1l1^-^V5^+^ cells near the infarct border (Fig. 3d). These results suggest that astrocytic specificity of the 4×6T cassette is largely maintained after a major CNS insult.

### *Astrocyte* specificity *of 4×6T cassette is preserved for long time periods and across CNS regions*

In the majority of experiments we evaluated astrocyte specificity two weeks post-viral delivery; while expression levels are robust at two weeks, they continue to increase^30^. Therefore, we assessed the stability of 4×6T astrocyte-specific expression over time. We systemically delivered PHP.eB::GfaABC1D-Cre-4×6T and PHP.eB::CAG-flex-lck-smV5-4×6T to young adult mice and assessed expression patterns six months later. Astrocyte specificity of 4×6T viral expression was maintained in the cortex at six months post-injection (99.87% ± 0.04 Sox9^+^; Fig 4a).

**Figure 4.**
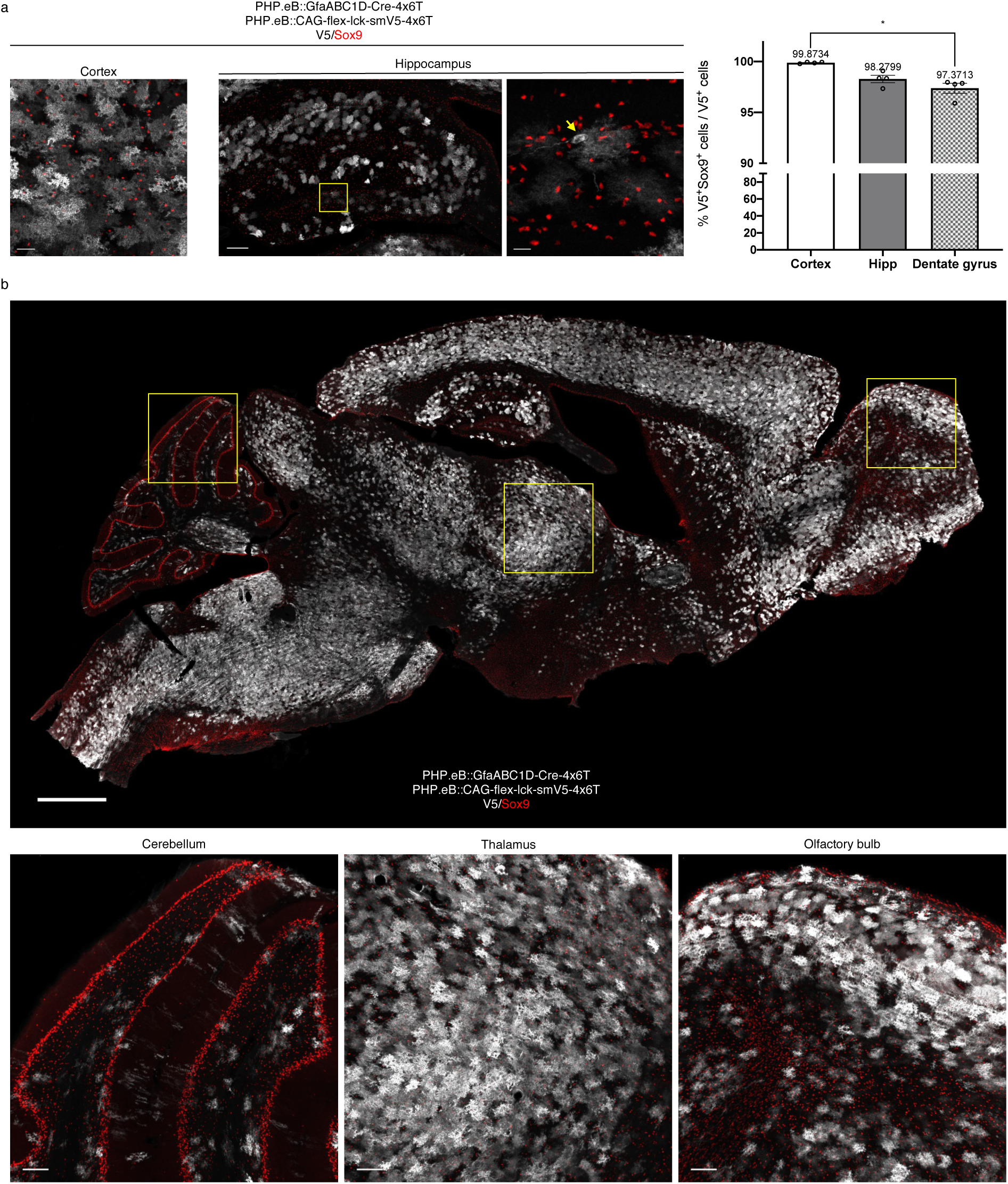
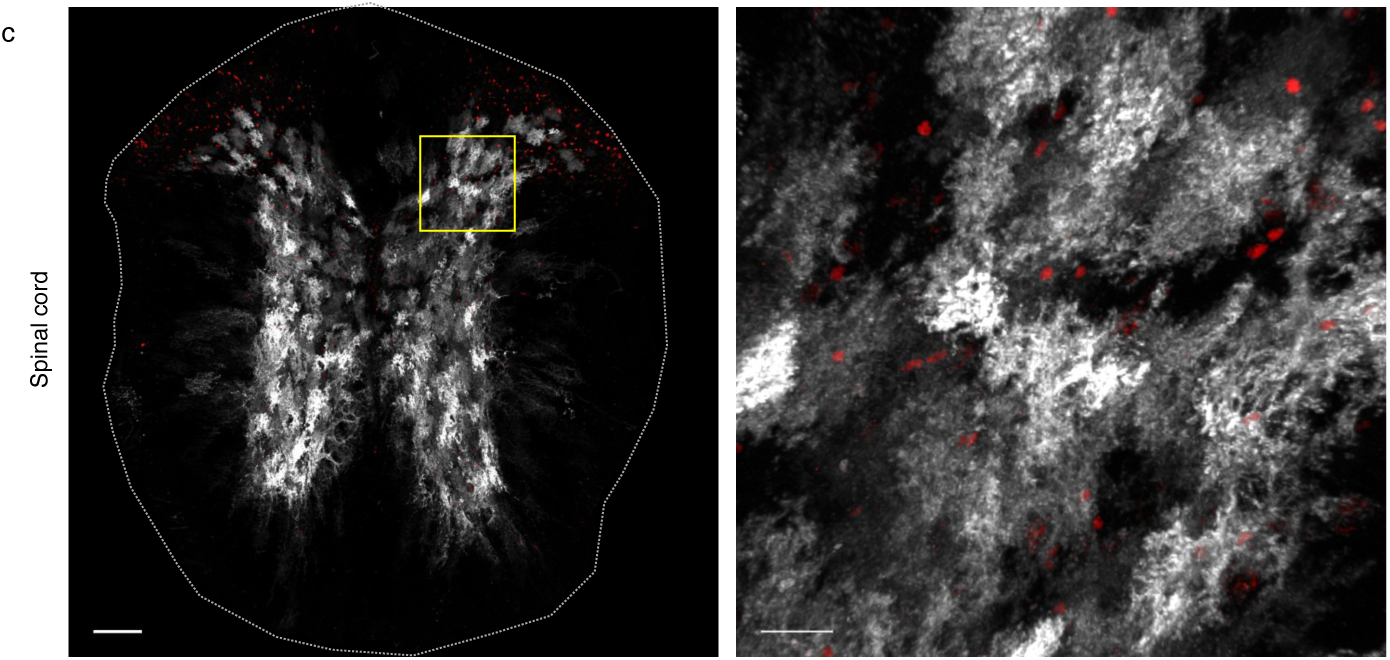
Astrocyte specificity of 4×6T cassette is preserved for long time periods and across CNS regions. **a,** Astrocyte specificity six months after injection (5×10^11 vg/mouse PHP.eB::GfaABC1D-Cre-4×6T; 5×10^11 vg/mouse PHP.eB::CAG-flex-lck-smV5-4×6T) into young adult (2-5 month old) mice is preserved in the cortex (scale bar: 40μm) and to a slightly lesser degree in the hippocampus. Hippocampus, left: entire structure (scale bar: 150μm); yellow box shows region of higher magnification on the right (scale bar: 20μm). Yellow arrow: example of neuron in the dentate gyrus. Astrocyte specificity is higher in the cortex than the dentate gyrus sub-region (*p=0.0134); Kruskal-Wallis test, Dunn’s multiple comparisons test, p=0.0024, Kruskal-Wallis statistic 8.346). Mean ± SEM; n=4 mice per brain region. **b**, Sagittal section of mouse brain six months after virus injection (scale bar: 1cm), with higher magnification examples of the cerebellum, thalamus, and olfactory bulb (scale bars: 100μm), showing high levels of colocalization of V5 with Sox9. **c**, Coronal section of mouse spinal cord six months after virus injection (scale bar: 100μm); yellow box denotes area of higher magnification section on the right (scale bar: 20μm), showing high V5/Sox9 colocalization.

Given the potential for transduced progenitor cells, we evaluated astrocyte specificity in the hippocampus. Astrocyte specificity remained high throughout the hippocampus (98.28% ± 0.36 Sox9^+^; Fig. 4a), although it was significantly lower than cortical specificity both in the hippocampus overall and in the dentate gyrus specifically (97.37% ± 0.49 Sox9^+^). Astrocyte specificity was high in the olfactory bulb, thalamus, cerebellum, and spinal cord (Fig. 4b, c); indeed, we could not identify a region in which there was overt non-astrocytic viral staining. Interestingly, white matter astrocytes in both the brain and spinal cord were largely resistant to systemic PHP.eB viral transduction.

### Transcriptomic *changes observed in astrocytes with viral transduction with and without 4×6T cassette*

One potential caveat to viral use is whether normal cellular processes are affected by viral transduction itself. Therefore, we evaluated astrocytic transcriptomes using RiboTag^31^ mice, a form of translating ribosomal affinity purification (TRAP). A requisite ribosomal subunit is tagged using Cre, allowing immunoprecipitation (IP) of ribosomes. Ribosomally loaded mRNA can then be purified and sequenced. We generated four groups of RiboTag astrocyte mice: intracortical or systemic delivery of PHP.eB::GfaABC1D-Cre-4×6T; intracortical delivery of AAV2/5::GfaABC1D-Cre, the current gold-standard approach for astrocyte-enriched viral manipulation; and transgenic RiboTag/Aldh1l1-CreERT2 mice^32^, to obtain astrocyte-specific labeling without viral transduction. These groups allow us to compare how viral transduction affects astrocytes; how the 4×6T cassette modifies those responses; and how the route of delivery impacts astrocytic transcriptomic responses.

We evaluated astrocyte specificity of RiboTag expression by immunostaining for the ribosomal tag (HA^+^, Fig. 5a). Aldh1l1-CreERT2 transgenic mice and systemic PHP.eB::GfaABC1D-Cre-4×6T were highly astrocyte-specific (Aldh1l1-CreERT2: 99.95% ± 0.03 Aldh1l1^+^; systemic Cre-4×6T: 99.96% ± 0.02 Aldh1l1^+^). Interestingly, both intracortical injections showed higher astrocyte specificity in Ribotag mice than in Ai14-tdTomato mice with similar injections (Fig 2d). As both transgenic lines are flox-stop-flox reporters, this may reflect differences in how easily Cre can induce recombination in different loci^33^. While we observed significant neuronal expression with AAV2/5::GfaABC1D-Cre (73.97% ± 2.90 Aldh1l1^+^), astrocyte specificity with intracortical PHP.eB::GfaABC1D-Cre-4×6T was high (98.79% ± 0.32 Aldh1l1^+^), reflecting enhanced astrocyte specificity of the 4×6T miR cassette. We also evaluated GFAP immunoreactivity, a rough metric of astrocyte reactivity^34^; intracortical injection with either serotype induced increases in GFAP, as expected, but systemic injection did not, suggesting that viral transduction alone does not induce overt reactivity (Fig. 5a).

**Figure 5.**
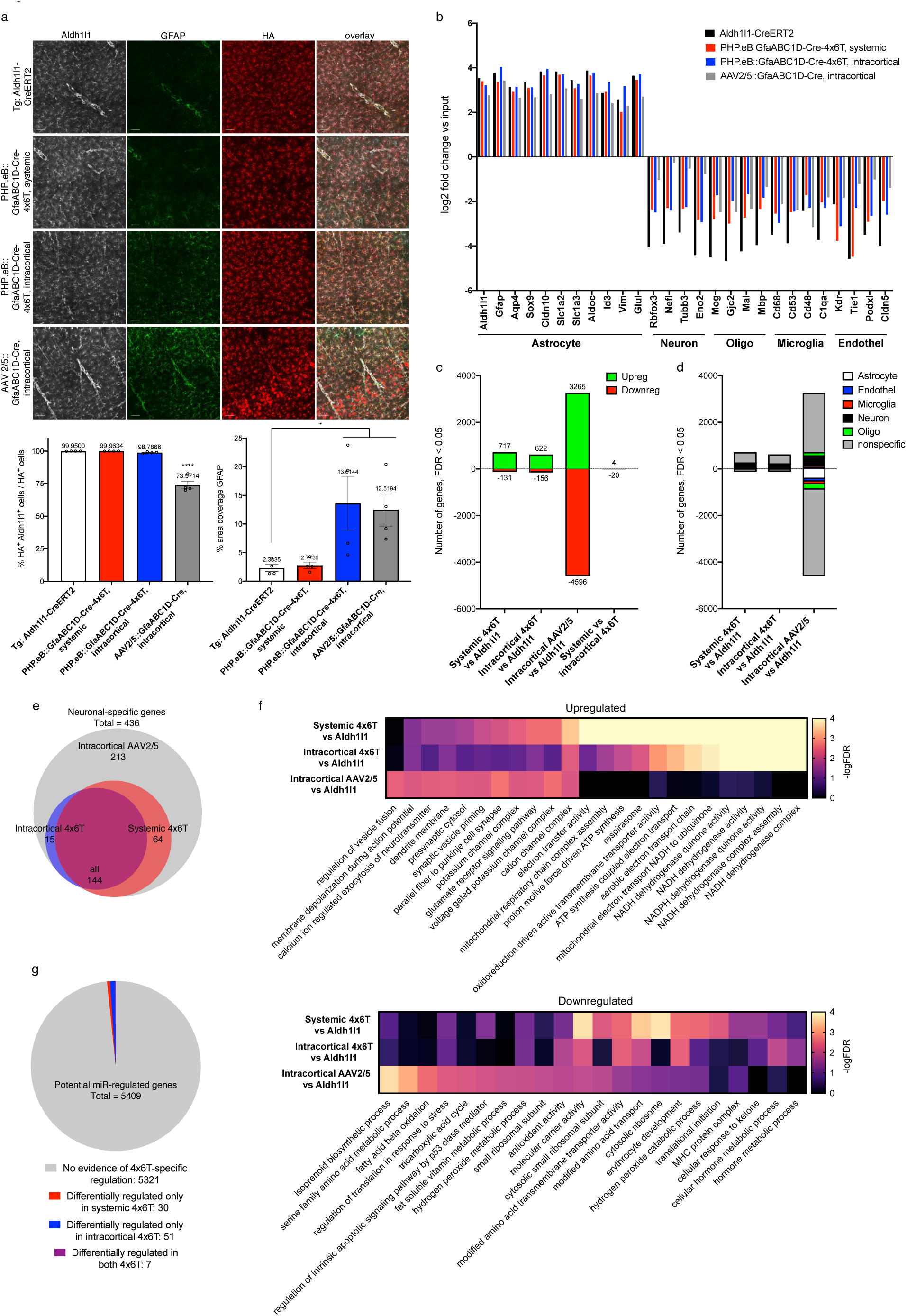
Transcriptional analysis of astrocytes with viral transduction with and without 4×6T cassette (TRAP). **a**, Immunohistochemical analysis of RiboTag^+^ cells (hemagglutinin HA^+^ ribosomal tag), colocalized with astrocytic marker Aldh1l1 and astrocytic reactivity marker GFAP. Astrocytic specificity is high in Aldh1l1-CreERT2 transgenic mice and systemic delivery of PHP.eB::GfaABC1D-Cre-4×6T with no overt evidence of astrocyte reactivity (GFAP % coverage). Specificity remains high with intracortical delivery of PHP.eB::GfaABC1D-Cre-4×6T and decreases with intracortical delivery of AAV2/5::GfaABC1D-Cre; this route of viral delivery shows some evidence of astrocyte reactivity by GFAP immunoreactivity. Scale bars: 50μm. Mean ± SEM; n=4 mice per cohort. Astrocyte specificity, HA^+^Aldh1l1^+^/HA^+^: one-way ANOVA, Holm-Sidak’s multiple comparisons test, p<0.0001, F=77.01, df=15; Aldh1l1-CreERT2 vs AAV2/5::GfaABC1D-Cre ****p<0.0001. GFAP % area coverage: Aldh1l1-CreERT2 = 2.33 % coverage ± 0.64; PHP.eB::GfaABC1D-Cre-4×6T systemic, 2.77 % coverage ± 0.55; PHP.eB::GfaABC1D-Cre-4×6T intracortical, 13.61 % coverage ± 4.71; AAV2/5::GfaABC1D-Cre, 12.52 % coverage ± 2.90; one-way ANOVA, Holm-Sidak’s multiple comparisons test, p=0.0011, F=10.54, df=15; Aldh1l1-CreERT2 vs PHP.eB::GfaABC1D-Cre-4×6T intracortical *p=0.0431; Aldh1l1-CreERT2 vs AAV2/5::GfaABC1D-Cre *p=0.0481. **b**, Relative levels of enrichment and de-enrichment of canonical genes for astrocytes, neurons, oligodendrocytes, microglia, and endothelial cells in IP vs input samples. **c**, Differentially expressed genes in IP samples: different viral cohorts vs Aldh1l1-CreERT2 IP samples, and systemic vs cortical 4×6T samples. FDR < 0.05, average FPKM across all IP samples > 1. **d**, Cell-type specificity of differentially expressed genes in IP samples: genes in (c) that are represented in the top 1000 specific genes for astrocytes, endothelial cells, microglia, neurons, or oligodendrocytes. **e**, Venn diagram showing the overlap of which neuron-specific genes from (d) are upregulated in different viral cohort IP samples vs Aldh1l1-CreERT2 samples. **f**, Top ten most significantly upregulated and downregulated Gene Ontology gene sets (FDR < 0.05) in ranked IP transcriptomes of each of the viral cohorts, calculated with Gene Set Enrichment Analysis; some gene sets overlapped, particularly for 4×6T cohorts, and only 7 gene sets passed FDR < 0.05 for intracortical 4×6T. In cases where the FDR was 0, scores were reassigned as 0.0001 to visualize the relative-logFDR. **g**, Numbers of genes potentially regulated by miRNAs that comprise the 4×6T cassette (5409 total) that show evidence of 4×6T-specific regulation in IP samples (FDR < 0.05 in either 4×6T cohort vs Aldh1l1-CreERT2 cohort; not differentially regulated in AAV2/5 cohort vs Aldh1l1-CreERT2 cohort). Note: none of the 5409 genes show evidence of 4×6T-specific regulation in input samples. n=4 mice per cohort.

TRAP transcriptomic data reflect the relative enrichment of ribosomally loaded mRNA vs whole-tissue, or input, mRNA. The effectiveness of the cell-type-specific immunoprecipitation, therefore, can be assessed by evaluating IP-vs-input enrichment of cell-type-specific genes and de-enrichment of genes associated with other cells. We found high astrocytic enrichment in Aldh1l1-CreERT2 mice and both 4×6T cohorts, with lower but still substantial astrocyte enrichment in AAV2/5 samples (Fig. 5b). De-enrichment of non-astrocytic genes was highest in Aldh1l1-CreERT2 mice, lower but substantial in both 4×6T cohorts, and lowest in the AAV2/5 cohort, reflecting the high levels of neuronal contamination in that cohort. Interestingly, endothelial decontamination was comparably robust in Aldh1l1-CreERT2 and both 4×6T cohorts, and weakest in AAV2/5 samples.

Comparing all IP experimental cohorts to Aldh1l1-CreERT2, we found the highest levels of differentially expressed genes (DEGs) in AAV2/5 (Fig. 5c): the greatest molecular difference in gene expression from Aldh1l1-CreERT2, a gold standard of astrocyte mRNA expression, occurs when only the AAV2/5 capsid and minimal GFAP promoter is used to restrict expression to astrocytes. Somewhat unexpectedly, there were few gene expression changes between both 4×6T cohorts (Fig. 5c), although GFAP was upregulated in the intracortical samples compared to systemic injection (Supp. Data 1), confirming our immunostaining findings. These results suggest that the route of viral administration has a minimal effect on astrocytic gene expression, with intracortical injection inducing only mild reactivity.

Given the lower de-enrichment of non-astrocytic genes in viral cohorts compared to Aldh1l1-CreERT2 (Fig. 5b), we explored which of the viral DEGs were cell-type-specific. We compared these gene lists to cell-type-specific gene signatures, which included the top 1000 most specific genes in astrocytes, endothelial cells, microglia, neurons, and oligodendrocytes, identified using three separate mouse brain cell transcriptomic studies^35^. Most DEGs were not overtly cell-type-specific; neurons made up the largest component of cell-type-specific upregulated genes (Fig. 5d). All of the neuronal upregulated genes in either 4×6T cohort were upregulated in AAV2/5 samples, while 48.8% of neuronal upregulated genes in AAV2/5 samples were not upregulated in either 4×6T cohort (Fig. 5e). The TRAP datasets reflect both any differences in which cell populations are labeled as well as virus-induced changes within the labeled cells: these cell-type-specific enrichment/de-enrichment and gene set analyses demonstrate the higher neuronal contamination of the AAV2/5 samples compared to 4×6T-modified virus, despite the fact that AAV2/5 is a more astrocyte-specific serotype.

To address which transcriptomic changes may occur in astrocytes due to virus transduction itself, we compared these IP datasets with datasets of astrocytes in related pathological settings as well as all Gene Ontology (GO) gene sets. Astrocytes are responsive to environmental perturbations, including pathologic viral infection and direct injury, of which direct intracortical injection is a mild form. The gene changes observed between Aldh1l1-CreERT2 astrocytes and AAV-transduced astrocytes may reflect common transcriptional pathways to those in astrocytes infected with pathologic viruses, to astrocytes after a stab wound injury, or to other biological processes listed in GO. To assess the physiological relevance of the transcriptional changes observed, we used Gene Set Enrichment Analysis^36^ (GSEA) to compare our datasets with transcriptomic changes in astrocytes treated with poly I::C to broadly mimic viral stimulation^37^, infected with Zika virus^38^ or HIV^39^, or had undergone a cortical stab wound^40^, and all GO gene sets to broadly assess these datasets. GSEA uses a ranked list of all transcriptional changes in the datasets of interest (AAV2/5 and 4×6T cohorts) and assesses where genes in a biologically defined test gene set fall along that ranked list.

Overrepresentation of a test gene set at either extreme of the dataset of interest suggests up- or down-regulation of that gene set and thus the biological pathway. None of the astrocyte-specific viral or injury datasets were significantly enriched in AAV2/5 or 4×6T-transduced astrocytes (FDR <0.05,Supp. Fig 1), suggesting that the gene changes observed in these cohorts do not reflect common transcriptional pathways induced with pathological viruses or stab wound. Numerous GO gene sets were significantly up- or down-regulated (AAV2/5: 305 up, 37 down; 4×6T intracortical: 40 up, 7 down; 4×6T systemic: 192 up, 12 down; Supp. Data 2); the most significantly upregulated pathways in AAV2/5 reflected neuronal expression, while the most significantly upregulated pathways in both 4×6T cohorts suggest changes in NADH activity (Fig 5f). The most significantly downregulated pathways in all cohorts are more varied but suggest that AAV transduction in astrocytes may impact metabolism and translation. Together, these results suggest that AAV transduction does not induce overt reactivity in astrocytes but may influence metabolic pathways.

One potential concern in using miRNA targeting sequences is whether those sequences might act as an miRNA sink, by binding endogenous miRNAs and thereby preventing their binding to and regulation of endogenous genes. Therefore, we generated a list of genes in the mouse genome that have shown interaction with the miRNAs in the 4×6T cassette, using the miRNA targeting database TarBase v8^41^ (5409 genes). We explored which of these genes showed evidence of differential regulation by 4×6T virus transduction both in input samples, which include neurons and endothelial cells that express the miRs, and in the astrocyte-enriched IP samples. No genes in the input samples were differentially regulated (FDR <0.05) in either 4×6T cohort compared to Aldh1l1-CreERT2. In IP samples, we evaluated changes that were specific to the presence of the 4×6T cassette: genes that were differentially regulated in either 4×6T cohort but not in the AAV2/5 cohort, reflecting potential altered expression due to interference with endogenous roles of miRs rather than due to AAV transduction. 88 of 5409 genes fit these criteria (Fig. 5g; Supp. Data 3), showing very little evidence for unintentional dysregulation of endogenous miRNA function due to the 4×6T cassette.

### Broader applicability of the 4×6T cassette in conferring astrocyte specificity

Cre is a particularly sensitive cargo protein to off-target expression, as very low Cre expression can induce genetic recombination. We wondered if the neuronal contamination issues seen with PHP.eB::GfaABC1D-Cre were present more broadly with PHP.eB::GfaABC1D vectors with less sensitive cargos and whether addition of the 4×6T might improve specificity. We systemically delivered a spaghetti monster reporter (smMyc; PHP.eB::GfaABC1D-lck-smMyc, 2×10^11 vg/mouse) to assess astrocyte specificity with a reporter protein and found high levels of non-astrocytic contamination (67.78% ± 2.98 Sox9^+^). In contrast, adding the 4×6T cassette (PHP.eB::GfaABC1D-lck-smMyc-4×6T) significantly improved specificity (99.80% ± 0.05 Sox9^+^) without affecting efficiency of astrocytic transduction (smMyc: 34.54% ± 0.97 Sox9^+^ cells transduced; smMyc-4×6T: 34.79% ± 0.55 Sox9^+^ cells transduced); in non-recombinase vectors, WPRE was included to enhance expression. We observed no difference in the strength of astrocytic Myc expression by immunohistochemistry.

We also generated smV5-4×6T and smFLAG-4×6T vectors; while smV5 showed similar expression patterns to smMyc, we found very sparse transduction with smFLAG when delivered systemically compared to the other smFPs. We found no difference in astrocyte specificity of smV5-4×6T delivered systemically vs direct intracortical injection (99.85% ± 0.01 Sox9^+^, systemic; 99.61% ± 0.17 Sox9^+^, intracortical; Fig. 6b). Further, we found that virtually all astrocytes within the core of the intracortically injected region were transduced; this high efficiency did not come with a trade-off in specificity. These experiments demonstrate that even non-recombinase cargo can show high levels of non-astrocytic expression, which can be improved with the 4×6T cassette.

**Figure 6.**
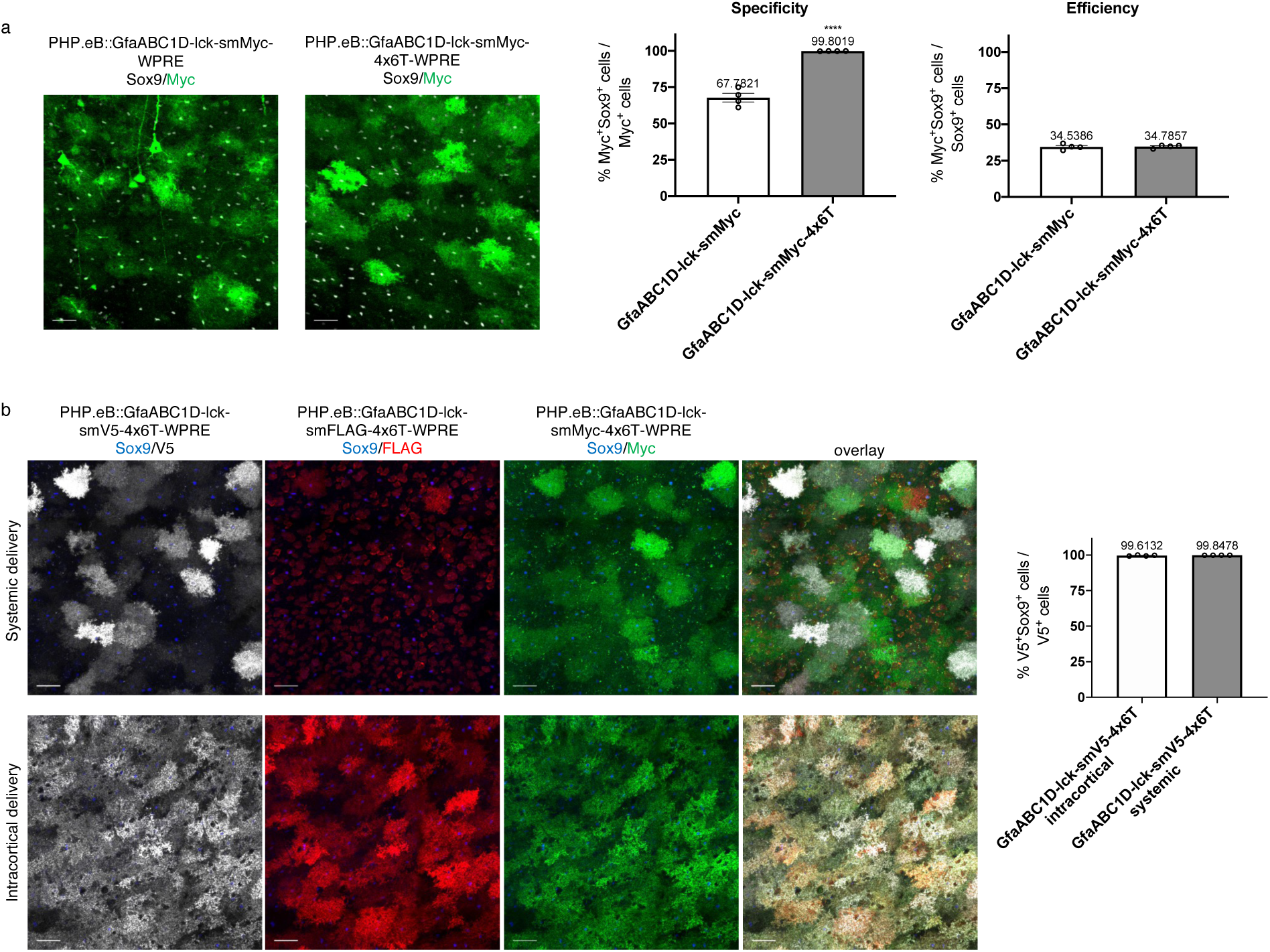
Non-recombinase astrocyte specificity with and without the 4×6T cassette. a, PHP.eB::GfaABC1D driving a spaghetti monster reporter (smMyc) shows high levels of non-astrocytic contamination, while adding a 4×6T cassette significantly increases astrocyte specificity (n=4 mice per cohort, two-tailed t-test, ****p<0.0001, t=10.73, df=6). The addition of the 4×6T cassette does not affect astrocyte transduction efficiency (two-tailed t-test, p=0.8314, t=0.2224, df=6). **b**, Astrocytic specificity remains high with spaghetti monster reporter smV5 constructs delivered either via direct intracortical injection or systemically (two-tailed t-test, p=0.2099, t=1.404, df=6). Adding the 4×6T cassette to other smFPs (FLAG, Myc) also show high astrocyte specificity, although PHP.eB::GfaABC1D-smFLAG-4×6T shows surprisingly low transduction efficiency when delivered systemically. Scale bars: 40μm. Note: all GfaABC1D-smFP constructs included a WPRE regulatory element to boost transgene expression.

One advantage of Cre is the development of inducible versions, including both tamoxifen-inducible (ERT2-Cre-ERT2^42^; ERCreER) and light-inducible (iCreV^43^) forms. To maximize the utility of Cre-based 4×6T viruses, we generated 4×6T versions of both ERCreER and iCreV. When we systemically delivered PHP.eB::GfaABC1D-ERCreER-4×6T or PHP.eB::GfaABC1D-iCreV-4×6T with PHP.eB::CAG-flex-lck-smV5, without including a 4×6T cassette on the flex reporter, we found extensive, predominantly neuronal contamination in the absence or presence of the inducing factor (tamoxifen or light, respectively) (Fig. 7a). By including 4×6T cassette on the reporter, however, this contamination was almost fully resolved, generating inducible astrocyte-specific viral manipulation, although we found higher astrocyte specificity with ERCreER (ERCreER, 99.95% ± 0.03 Sox9^+^; iCreV, 97.75% ± 0.52 Sox9^+^; Fig. 7a).

**Figure 7.**
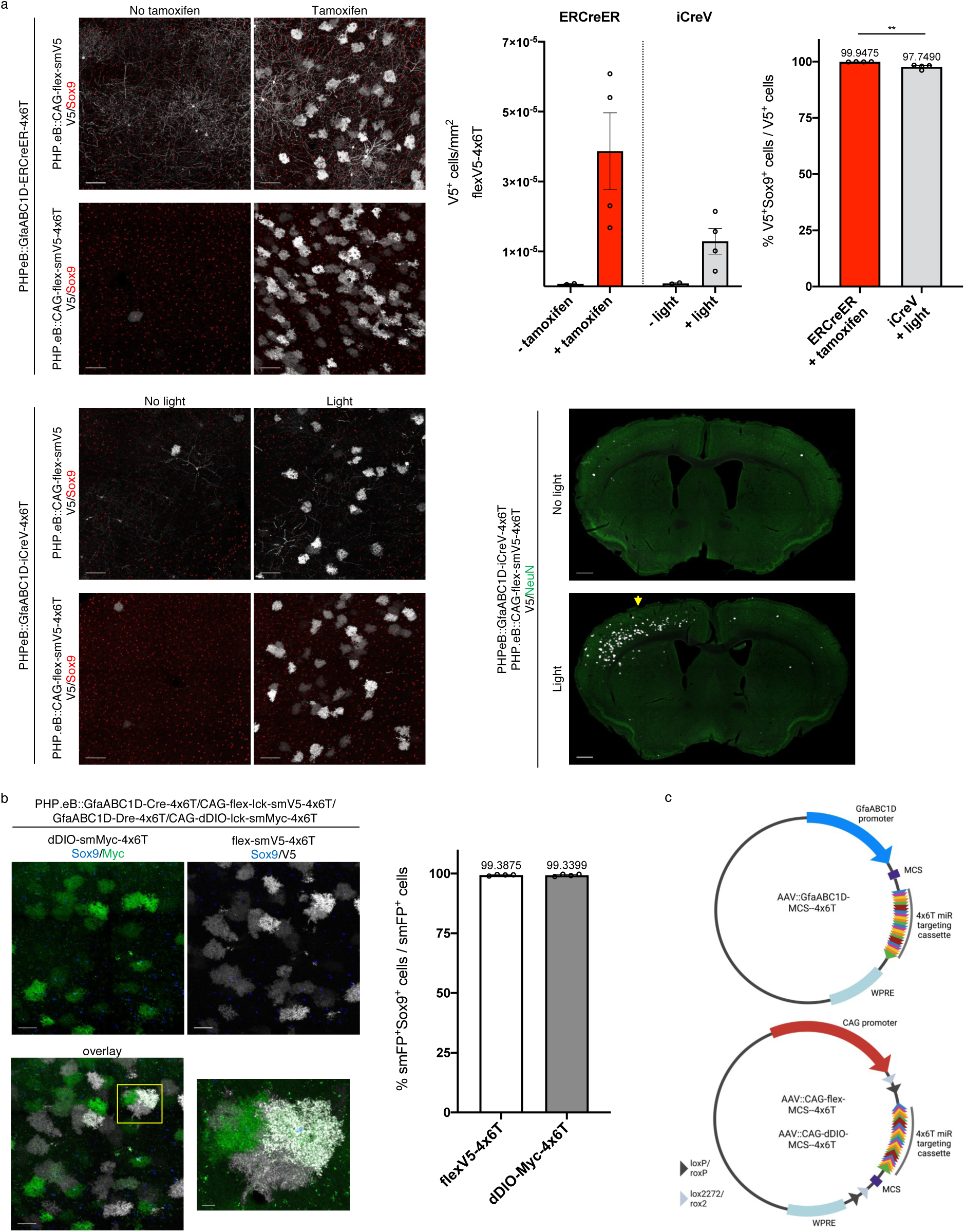
Astrocyte specificity of the 4×6T cassette with inducible and alternative recombinases. **a**, Tamoxifen-(ERCreER) and light-(iCreV) inducible forms of Cre show high levels of neuronal background without induction when co-injected with a flexV5 reporter with no 4×6T cassette (PHP.eB::CAG-flex-smV5) but much less background when co-injected with a flexV5-4×6T reporter (PHP.eB::CAG-flex-smV5-4×6T). Scale bars: 100μm. Astrocyte specificity with a flexV5-4×6T reporter is high with both inducible forms of Cre, but higher in ERCreER (two-tailed unpaired t-test, **p=0.0055, t=4.233, df=6; n=4 mice with tamoxifen or light; 2 mice with no tamoxifen or no light). Light induction of iCreV shows expression across a broad area of cortex ipsilateral to light placement and limited contralateral expression. Light placement: yellow arrow. Scale bar: 500μm. **b**, PHP.eB::GfaABC1D-Dre-4×6T and Dre-dependent reporter PHP.eB::CAG-dDIO-smMyc-4×6T can be used orthogonally with Cre/flex viral systems and show similar levels of astrocytic specificity (scale bar: 50μm); yellow box shows region of higher magnification on the right (scale bar: 10μm); n=4 mice per recombinase. All data presented as mean ± SEM. **c**, Schematic diagrams of empty vectors with multiple cloning sequences (MCS) for insertion of other cargo: GfaABC1D-MCS--4×6T for non-recombinase-dependent expression; CAG-flex-MCS--4×6T for Cre-dependent expression; and CAG-dDIO-MCS--4×6T for Dre-dependent expression. Note the presence of a woodchuck hepatitis virus post-transcriptional regulatory element (WPRE); while we omitted this element in the case of recombinase vectors, where high levels of transgene expression were neither wanted nor needed, we have included it in these more general vectors. More detail on transgene components and the full sequences can be found on Addgene.

In order to make an astrocyte-specific DNA recombinase that could be used orthogonally with Cre, we generated GfaABC1D-Dre-4×6T and a Dre-dependent reporter, CAG-dDIO-lck-smMyc-4×6T^44^. We co-injected PHP.eB::GfaABC1D-Cre-4×6T, PHP.eB::CAG-flex-lck-smV5-4×6T, PHP.eB::GfaABC1D-Dre-4×6T, and PHP.eB::CAG-dDIO-lck-smMyc-4×6T, and found similar astrocyte specificity with both recombinase/reporter systems (Cre, 99.39% ± 0.15 Sox9^+^; Dre, 99.34% ± 0.23 Sox9^+^; Fig. 7b). At non-saturating titers there is some, but not complete, overlap in Dre-and Cre-labeled cells; these viruses, therefore, can be used to label neighboring astrocytes with different combinations of reporters to evaluate astrocytic tiling (Fig. 7b).

The 4×6T vectors presented here – GfaABC1D-Cre, ERCreER, iCreV, Dre, and smFPs; conditional reporters CAG-flex-lck-smV5 and dDIO-lck-smMyc – are ready-to-package AAV plasmids for astrocyte-specific manipulation and morphologic analysis (Table 1). In order to facilitate the addition of 4×6T to other cargo as needed, we further generated empty vectors in which the cargo protein was replaced with a multiple cloning sequence (MCS). We generated three such vectors: GfaABC1D-MCS--4×6T, CAG-flex-MCS--4×6T, and CAG-dDIO-MCS--4×6T (Fig. 7c; Table 1). Together, these vectors form a toolbox of astrocyte-specific AAV plasmids to facilitate astrocytic viral manipulation across a wide range of experimental conditions.

**Table 1.**
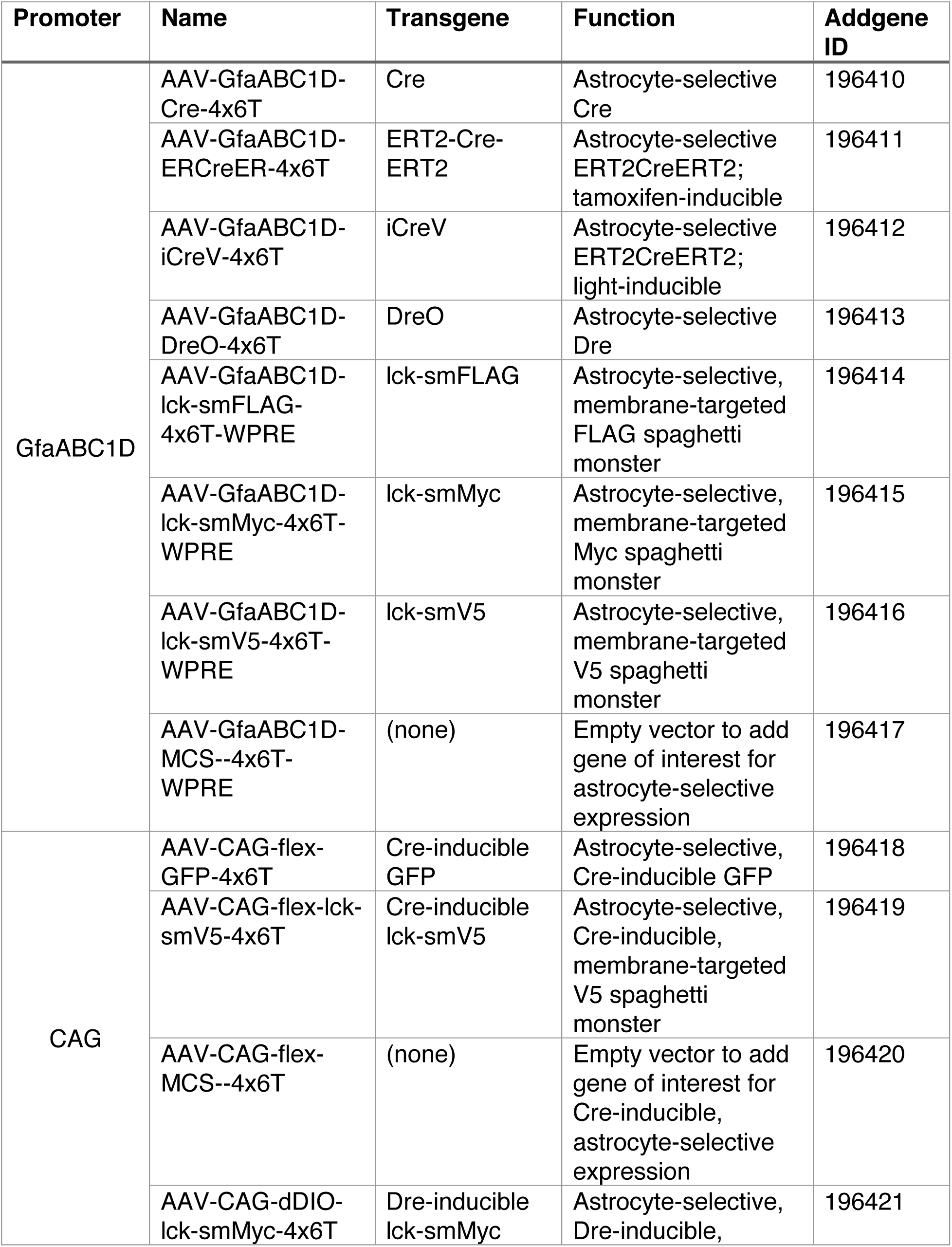
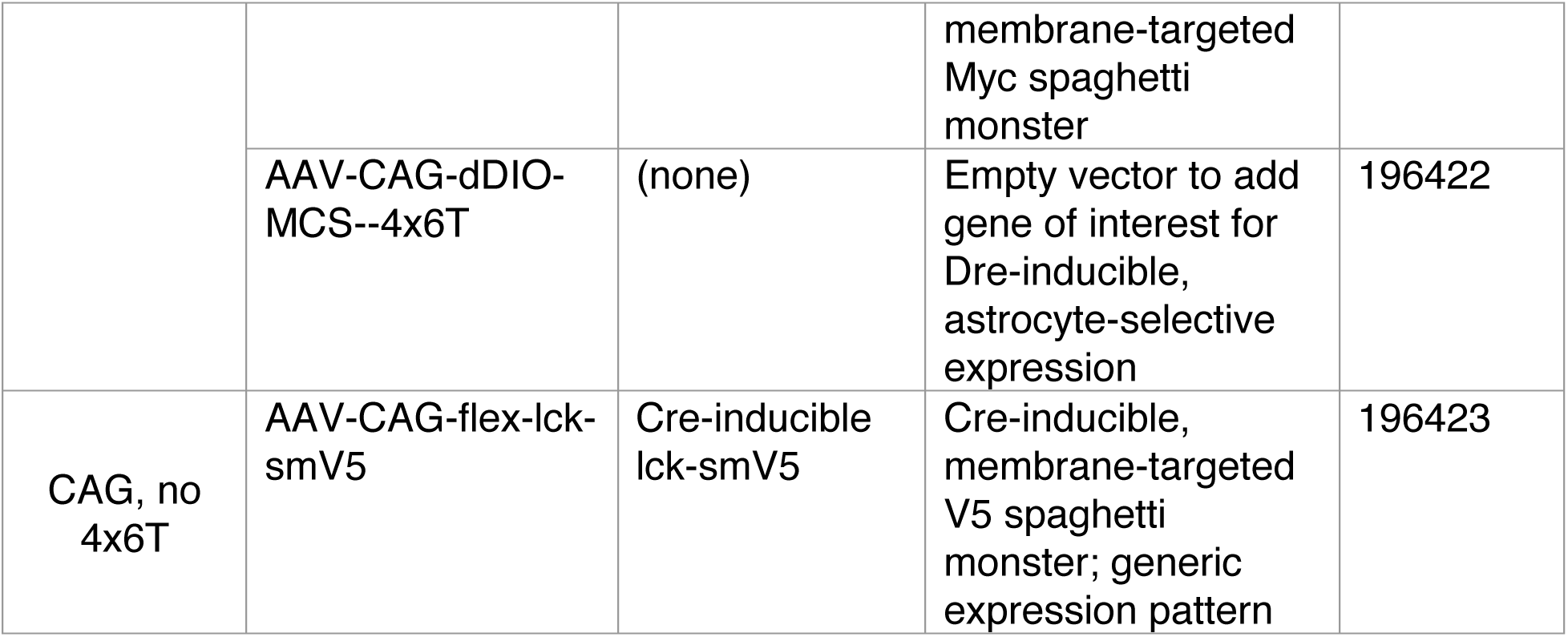
Plasmids deposited at Addgene.

## Discussion

Here, we present the development of a miRNA targeting cassette comprised of four copies of each of six miR targeting sequences (4×6T). This multiplexed cassette de-targets both neurons and endothelial cells, demonstrating the flexible use of miR targeting in improving viral specificity by decreasing multiple off-target populations simultaneously. The 4×6T cassette (i) confers a high level of astrocyte specificity on AAV vectors across all serotypes and cargos tested, (ii) exhibits specificity across the mouse lifespan and in injury conditions, and (iii) maintains astrocyte-specific expression patterns for at least 6 months post-injection. We have used the 4×6T cassette to create a toolbox of astrocytic AAV vectors that will enable cleaner, more flexible astrocytic genetic manipulation across a wide range of experimental conditions.

While we focused this analysis on the cerebral cortex, we found that systemic delivery of PHP.eB::GfaABC1D viruses transduces astrocytes across the CNS, with the notable exception of white matter, where we found very low transduction. White matter astrocytes were transduced with intracortical injection of different serotypes, suggesting this may be an issue of serotype and/or delivery route. Although the 4×6T cassette dramatically improved astrocyte specificity of all tested serotypes, given the level of neuronal contamination upon intracortical high-titer injection of PHP.eB::CAG-flex-lck-smV5-4×6T, it appears that high titer combined with a generic promoter can overcome miR regulation. Therefore, for high titer direct injections of flex cassettes with a strong generic promoter, we recommend AAV2/5 for the highest possible astrocytic specificity. We also found slightly lower astrocytic specificity in the hippocampus, particularly the dentate gyrus; however, even when high-titer virus was delivered to adult mice the level of off-target labeling remained quite low, suggesting that there may be some de-targeting of progenitor cells by the 4×6T cassette. This would be consistent with the high miR124 levels found in progenitor cells^45^.

This modification to improve astrocyte specificity is compatible with different capsids, makes it possible to take advantage of the growing capabilities of designer AAV capsids; here, we used the systemically deliverable PHP.eB capsid, but 4×6T could be used with serotypes with other capabilities, such as delivery to the peripheral nervous system (PHP.S^4^). The 4×6T cassette is also promoter-independent and could be used with different promoters or enhancers to potentially restrict expression to specific astrocytic subpopulations. Given the extensive and growing evidence of astrocyte heterogeneity^46–49^, the ability to rapidly, specifically genetically manipulate distinct astrocytic subpopulations would be extremely powerful; the 4×6T cassette makes that more feasible and opens up new avenues for astrocyte-specific gene therapy.

The repeating nature of the cassette makes *de novo* synthesis challenging and complicates PCR-based cloning. In contrast, the empty vectors presented here facilitate rapid, easy generation of 4×6T vectors with any number of cargo. These vectors, combined with the reporters and recombinases generated here, provide a wide range of options to enable highly specific, rapid genetic manipulation of astrocytes.

## Methods

### Generation of miR-containing plasmids

Plasmids were constructed using Gibson assembly^50^. Viral backbones were generated by restriction enzyme digest. Inserts were generated by polymerase chain reaction (PCR); primers were designed using NEBuilder (New England BioLabs). All plasmids were confirmed by whole-plasmid sequencing (Primordium Labs).

AAV::GfaABC1D plasmids: AAV::GfaABC1D plasmids were based on backbone plasmid AAV-pmSyn1-EBFP-Cre^51^. AAV pmSyn1-EBFP-Cre was a gift from Hongkui Zeng (Addgene plasmid # 51507; http://n2t.net/addgene:51507; RRID:Addgene_51507). Two copies of single miR-T sequences were added in primers, split between the 3’ end of the Cre insert and the 5’ end of hGH polyA regulatory sequence insert. The 4×6T cassette was constructed in multiple steps, which was necessary due to complexities in synthesizing high numbers of repeating DNA sequences. Two copies of each of the 5 neuronal miR-T (2×5T) were synthesized as a single gene product (gBlock, Integrated DNA Technologies) and added to a Cre plasmid. This 2×5T plasmid was then used as part of the backbone to add an additional two copies of each neuronal miR via PCR insert, to which four copies of miR-126T were added in primers. Additional restriction enzyme sites were added in primers to facilitate downstream cloning. AAV::GfaABC1D-Cre-4×6T plasmid was used as a backbone to generate other GfaABC1D-(cargo)-4×6T plasmids. ERCreER: pCAG-ERT2CreERT2 was a gift from Connie Cepko (Addgene plasmid # 13777; http://n2t.net/addgene:13777; RRID:Addgene_13777)^42^. iCreV: pAAV-EF1a-iCreV was a gift from Hongkui Zeng (Addgene plasmid # 140135; http://n2t.net/addgene:140135; RRID:Addgene_140135)^43^. Dre: AAV phSyn1(S)-DreO-bGHpA was a gift from Hongkui Zeng (Addgene plasmid # 50363; http://n2t.net/addgene:50363; RRID:Addgene_50363)^51^. In GfaABC1D-smFP-4×6T-WPRE plasmids, WPRE was added to boost spaghetti monster fusion protein (smFP) reporter expression. pCAG_smFP V5 (Addgene plasmid # 59758; http://n2t.net/addgene:59758; RRID:Addgene_59758), pCAG_smFP FLAG (Addgene plasmid # 59756; http://n2t.net/addgene:59756; RRID:Addgene_59756), and pCAG_smFP Myc (Addgene plasmid # 59757; http://n2t.net/addgene:59757; RRID:Addgene_59757) were gifts from Loren Looger^23^.

AAV::CAG plasmids: AAV::CAG plasmids were based on backbone plasmid AAV-FLEX-GFP. pAAV-FLEX-GFP was a gift from Edward Boyden (Addgene plasmid # 28304; http://n2t.net/addgene:28304; RRID:Addgene_28304). Reverse-complement inserts for smFPs and the 4×6T cassette were generated via PCR; given the repeating nature of the 4×6T cassette, multiple 4×6T PCR products were generated. These products were separated using 3% low-melting-temperature agarose gels to ensure isolation and purification of the proper molecular weight product. Flex construct lox sites were replaced with rox sites to generate Dre-dependent dDIO constructs, using pAAV-Ef1a-dDIO hChR2(H134R)-EYFP; pAAV-Ef1a-dDIO hChR2(H134R)-EYFP was a gift from Karl Deisseroth (Addgene plasmid # 55640; http://n2t.net/addgene:55640; RRID:Addgene_55640)^44^.

Empty vector plasmids: Existing GfaABC1D-smFP-4×6T-WPRE and CAG-flex/dDIO-smFP-4×6T plasmids were used as backbone plasmids, with cargo removed via restriction enzyme digest. Multiple cloning sequences (MCS) comprised of restriction enzyme sites that were absent from the backbone plasmid were generated via gene synthesis (Integrated DNA Technologies) and added via Gibson assembly.

### AAV packaging and titration

All AAVs used in this study were produced in the Carmichael lab, in accordance with the US National Institutes of Health Guidelines for Research Involving Recombinant DNA Molecules and the University of California Los Angeles Institutional Biosafety Committee, with the exception of PHP.eB::GFAP-Cre. pAAV.GFAP.Cre.WPRE.hGH was a gift from James M. Wilson (Addgene viral prep # 105550-PHPeB; http://n2t.net/addgene:105550; RRID:Addgene_105550).

Rep/cap plasmids: PHP.eB: pUCmini-iCAP-PHP.eB was a gift from Viviana Gradinaru (Addgene plasmid # 103005; http://n2t.net/addgene:103005; RRID:Addgene_103005)^4^. AAV2/1: pAAV2/1 was a gift from James M. Wilson (Addgene plasmid # 112862; http://n2t.net/addgene:112862; RRID:Addgene_112862). AAV2/5: pAAV2/5 was a gift from Melina Fan (Addgene plasmid # 104964; http://n2t.net/addgene:104964; RRID:Addgene_104964). AAV2/9: pAAV2/9n was a gift from James M. Wilson (Addgene plasmid # 112865; http://n2t.net/addgene:112865; RRID:Addgene_112865).

AAV packaging and titration were performed as described^52^, with the following modifications:

Step 27: Amicon filtration columns were covered with 15ml DPBS + 0.1% Pluronic F-68 for 10 minutes; solution was discarded and replaced with 15ml DPBS + 0.01% Pluronic F-68. Columns were centrifuged at 3000*g for 3 minutes; flow-through was discarded. DPBS + 0.001% Pluronic F-68 was used in place of DPBS for steps 27-30.

Step 39: qPCR primers were used based on transfer plasmid; WPRE primers used were as described^52^. hGH primers: Fwd: CCT GGG TTC AAG CGA TTC T. Rev: CAG CCT GGC CAA TAT GGT.

### Animals

Animal procedures were performed in accordance with the US National Institutes of Health Animal Protection Guidelines and the University of California Los Angeles Chancellor’s Animal Research Committee. All mice were housed in a facility with 12hr-12hr light-dark cycle, controlled temperature and humidity, and had ad libitum access to food and water. Young adult (8-10 week) C57Bl/6J mice were purchased from Jackson Labs (000664) and injected between 2 and 5 months of age; aged C57Bl/6J mice (18-20 month) were obtained from the National Institute on Aging Aged Rodent Colony and injected at 28 months of age. Conditional Cre-dependent mouse lines used were Ai14 (Rosa-CAG-LSL-tdTomato-WPRE)^19^ as a general reporter and RiboTag (loxP-STOP-loxP-Rpl22-HA)^31^ for TRAP transcriptomic analysis. RiboTag mice were crossed with Aldh1l1-Cre/ERT2^32^ to generate an astrocyte-specific TRAP line; Cre-negative, RiboTag-positive littermates were used for viral delivery TRAP. Three to five mice were used per condition for each experiment and included both male and female mice.

### AAV delivery, Cre induction, and stroke

Systemic AAV delivery in adult mice was performed via retroorbital injection, as described^52^. Virus was diluted in sterile saline to a standard volume of 50μl per mouse; as needed based on titer, volume was increased up to 100μl. Systemic AAV delivery in P1-2 pups was performed via temporal vein as described^53^, in a final volume of 25μl per animal, diluted in sterile saline. Systemic virus was delivered at 5×10^11 vg/mouse per virus, with the following exemptions: GFAP-Cre, 5×10^10 vg/mouse (Fig 1a); high titer, 3×10^12 vg/mouse (Fig. 2a); P1-2 pups, PHP.eB::GfaABC1D-Cre-4×6T + 2×10^10 vg/mouse PHP.eB::CAG-flex-lck-smV5-4×6T (Fig. 3a, b); 28 month old mice, 2×10^11 vg/mouse (Fig. 3a); systemic PHP.eB::GfaABC1D-Cre-4×6T in Ribotag mice, 1×10^12 vg/mouse (Fig 5a); PHP.eB:::GfaABC1D-lck-smMyc-WPRE, PHP.eB::GfaABC1D-lck-smMyc-4×6T-WPRE, PHP.eB::GfaABC1D-lck-smFLAG-4×6T-WPRE, PHP.eB::GfaABC1D-lck-smV5-4×6T-WPRE, 2×10^11 vg/mouse (Fig 6a, b); PHP.eB::GfaABC1D-Cre-4×6T/CAG-flex-lck-smV5-4×6T/CAG-dDIO-lck-smMyc-4×6T: 1×10^11 vg/mouse per virus, GfaABC1D-Dre-4×6T: 2×10^11 vg/mouse (Fig 6c).

Intracortical virus was delivered under isoflurane anesthesia in a stereotactic apparatus via pulled glass micropipette. 500nl of virus diluted in sterile saline was delivered to each of two locations in the cortex (A/P: 0.0mm, M/L: 2.0mm, D/V: −0.75mm; A/P: −1.2mm, M/L: 2.0mm, D/V: −0.65mm). For RiboTag intracortical cohorts, a third injection was performed in contralateral cortex (A/P: −0.6mm, M/L: −2.0mm, D/V: −0.75mm); ipsilateral tissue was used for ribosome immunoprecipitation, while contralateral tissue was used for immunohistochemistry. The pipette was left in situ for 3 minutes to allow proper virus diffusion. Intracortical virus was delivered at 1×10^12vg/ml in all experiments.

Cre induction: Tamoxifen was administered at 100mg/kg in corn oil i.p. for five days, beginning two weeks after virus injection (PHP.eB::GfaABC1D-ERCreER-4×6T) or on the day of virus administration in other RiboTag experimental cohorts (Aldh1l1-CreERT2 transgenic mice). Light induction: Two weeks after virus administration, (PHP.eB::GfaABC1D-iCreV-4×6T) mice were placed in a stereotactic apparatus under isoflurane anesthesia. A cold light source (KL1500 LCD; Carl Zeiss MicroImaging) attached to a 40x objective was positioned at the skull (A/P: 0.0mm, M/L: 2.0mm) and illuminated at maximum illumination for 40 minutes.

Middle cerebral artery occlusion: Distal middle cerebral artery occlusion was produced as previously described^54^. Briefly, the distal branch of the middle cerebral artery was permanently occluded via electrocoagulation, followed by 15 minute bilateral jugular vein clamp. Rectal temperature was maintained at 37°C ± 0.5°C throughout surgery.

### Tissue processing and immunohistochemistry

Animals were euthanized two weeks post-injection or post-Cre-induction; for long-term expression, animals were euthanized six months post-injection. Animals were exposed to terminal levels of isoflurane and transcardially perfused with PBS followed by cold 4% paraformaldehyde in PBS (PFA). The brain was removed, postfixed in 4% PFA for 3-6 hours, and dehydrated in 30% sucrose in PBS for two days before being snap frozen and stored at −80°C until sectioning. Brains were sectioned at 50μm on a cryostat (Leica Biosystems).

Fluorescence immunohistochemistry was performed on fixed frozen 50μm tissue sections. Briefly, sections were washed 3×PBS, underwent antigen retrieval (10mM sodium citrate, pH6, 80°C for 30min), washed 3×PBS, and blocked with 5% normal donkey serum (NDS) and 0.3% TritonX-100 in PBS for one hour. In experiments using biotin-labeled secondary antibody, NDS block was followed by avidin/biotin block: 0.001% avidin in PBS for 15min, 2×PBS washes, 0.001% biotin in PBS for 15min, 2×PBS washes. Sections were incubated in primary antibody with 2% NDS/0.3% TritonX-100 in PBS for at least 48 hours at 4°C. Sections were washed 3×PBS + 0.3% TritonX-100 and incubated in secondary antibody 1-2 hr at room temperature, followed by 3×PBS washes. In all experiments using flex-GFP virus, GFP fluorescence was further amplified with an anti-GFP 488-conjugated nanobody. Experiments using biotin-labeled secondary included an additional 1hr incubation in VioBlue-streptavidin. Labeled primary antibodies were applied after secondary and incubated for at least 48 hours at 4°C. Sections were mounted on gelatinized slides, dehydrated in ascending ethanol washes, cleared in 2× xylene washes, and coverslipped for imaging. Primary antibodies used in this study: GFAP (1:1000, chicken, Rockland #200-901-D60); Aldh1l1 (1:400, rabbit, Abcam #Ab87117); Sox9 (1:1000, rabbit, Millipore Sigma #AB5535); NeuN (1:1000, chicken, Synaptic Systems #266 006); CD31 (1:100, rat, BD Biosciences #550274); V5 (1:400, human, Absolute Antibodies #AB00136-10.0); Myc (1:400, 488-labeled, Biotium #20436); FLAG (1:400, 543-labeled, Biotium #20433); HA (1:100, rat, Roche #11-867-423), and GFP (1:500, 488-labeled nanobody, ChromoTek #GBA488-100). Secondary antibodies: 1:1000, Jackson ImmunoResearch, donkey anti-rabbit (488: 711-546-152; Cy3: 711-166-152; 647: 711-606-152), donkey anti-chicken (488: 703-546-155; Cy3: 703-166-155); donkey anti-rat (Cy3: 712-166-153; 647: 712-606-153); donkey anti-human (488: 709-546-159; 647: 709-606-149); 1:250, VioBlue-streptavidin (Miltenyi Biotec, 130-106-933).

Images were analyzed using Imaris software (Oxford Instruments, v9.9) to identify and colocalize virally transduced cells. For the majority of systemic injection experiments, analysis was restricted to the cortex and included at least two sagittal sections and 2.5mm anterior-posterior spread; for light-inducible iCreV and stroke experiments, samples were sectioned coronally to allow comparison to the non-induced/non-lesioned hemisphere. Direct intracortical virus injection tissue was sectioned coronally; analysis was restricted to the cortex and the core of the injected area and included at least two sections. GFAP percentage area covered was analyzed using Fiji Is Just ImageJ^55^. Briefly, a constant threshold was applied to maximum intensity projection images. The area of HA^+^ RiboTag cells in cortex was outlined as a region of interest (ROI), and this outlined ROI was applied to the GFAP channel. Thresholded GFAP^+^ area as a percentage of ROI was measured. Analysis included two contralateral sections from each of four animals per RiboTag cohort; ipsilateral tissue was used in transcriptomic analysis.

### Ribosomally-loaded mRNA isolation, sequencing, and analysis

Animals were exposed to terminal levels of isofluorane and transcardially perfused with 20mL cold PBS. The brain was rapidly removed, placed in a pre-chilled coronal brain matrix (Zivic Instruments) on ice, and divided into left and right hemispheres. The right hemisphere was placed in 4% PFA for 24 hours and processed for immunohistochemistry. The left hemisphere was further divided into 1mm coronal sections in the brain matrix. Relevant brain slices based on viral spread patterns (A/P +0.6mm to −1.8mm) were transferred to a dissection microscope in cold PBS on ice and the cortex was dissected (∼M/L 1.0mm to 3.0mm). Tissue was transferred to 1.5ml RNase/DNase-free tubes, weighed, snap frozen, and stored at −80°C. Immunoprecipitations were performed as described^56^; all samples were processed in parallel to minimize batch effects. RNA was extracted using the RNeasy Micro Plus kit (Qiagen).

Immunoprecipitated pull-down RNA and the corresponding input RNA were examined on RNA Pico chips (Agilent) before use. All of RNA Integrity Numbers (RIN) are above 8. cDNA libraries were prepared with SMART-Seq v4 RNA Ultra Low Input (Takara) + Nextera XT (Illumina). Paired-end 100bp sequences were generated over 1 lane by NovaSeq6000 using S4FC (Illumina).

After demultiplexing samples, we obtained between 73 and 109 million reads per sample (average: 88M). Quality control was performed on base qualities and nucleotide composition of sequences. Alignment to the M. musculus (mm10) refSeq (refFlat) reference gene annotation was performed using the STAR (v2.7.5c) with default parameters. Additional QC was performed after the alignment using PicardTools. Average uniquely mapped read rate was 86.2±0.9(SD)%. Total counts of read-fragments aligned to candidate gene regions were derived using HTSeq program (v0.12.4) and used as a basis for the quantification of gene expression.

Genes with CPM > 0.25 in at least 3 samples were selected for this analysis. Differential expression analysis was performed using the Limma-voom Bioconductor package (v3.50.1) and differentially expressed genes were selected based on FDR < 0.05 (false discovery rate, Benjamini Hochberg-adjusted p values). Paired-analysis mode was used for IP vs Input. To evaluate cell-type-specific IP vs input enrichment, cell-type specific genes were identified for astrocytes, neurons, endothelial cells, microglia, and oligodendrocytes^46,57^. To evaluate genes that were differentially expressed between IP samples from viral cohorts vs Aldh1l1-CreER mice, genes were filtered for average FPKM>1 in all IP samples and those with an FDR < 0.05 were extracted. Up- and down-regulated gene lists from each viral cohort were evaluated for representation in the top 1000 most specific genes in astrocytes, endothelial cells, microglia, neurons, and oligodendrocytes using mouse transcriptomic datasets^35^; overlap among upregulated neuronal genes across cohorts was assessed and visualized using BioVenn^58^. For Gene set Enrichment Analysis (GSEA), IP viral cohort vs Aldh1l1-CreER genes were sorted by (log2FC*-log10 pVal) to incorporate both the magnitude of the change and the statistical significance and then used as input for a preranked weighted analysis^36^. Reference gene sets were obtained for virally infected astrocytes^37–39^ and stab wound astrocytes^40^ and from the Molecular Signatures Database^59^ for GO and were filtered for gene sets with FDR < 0.05. For visualization purposes, gene sets with an FDR = 0 were reassigned a score of 0.0001, reflecting a value lower than the lowest non-0 score. Potential miRNA target genes were identified using TarBase v.8^41^; all *Mus musculus* target genes for each miR in the 4×6T cassette (miR124, miR126, miR137, miR329, miR369, miR431) were compiled and used to filter the complete gene list of both input and IP samples. For both IP and input, genes differentially regulated between either 4×6T cohort and Aldh1l1-CreER samples but not differentially regulated between AAV2/5 and Aldh1l1-CreER samples were extracted.

### Statistics and reproducibility

Statistical analysis of repeated measures were performed in Prism 8 (GraphPad). Data were subjected to normality testing (Shapiro-Wilk test); all data except figure 4a passed normality test and were analyzed using two-tailed t-tests or one-way ANOVAs with Tukey’s or Holm-Sidak’s multiple comparisons tests. Figure 4a failed normality and were analyzed using the nonparametric Kruskal-Wallis test with Dunn’s multiple comparisons test. P values < 0.05 were considered statistically significant and are reported in figure legends. All immunohistochemistry analyses were repeated in at least three mice and represent at least two sections per mouse spread along the viral transduction axis.

## Data availability

The following plasmids are available on Addgene: AAV-GfaABC1D-Cre-4×6T (196410), AAV-GfaABC1D-ERCreER-4×6T (196411), AAV-GfaABC1D-iCreV-4×6T (196412), AAV-GfaABC1D-DreO-4×6T (196413), AAV-GfaABC1D-lck-smFLAG-4×6T-WPRE (196414), AAV-GfaABC1D-lck-smMyc-4×6T-WPRE (196415), AAV-GfaABC1D-lck-smV5-4×6T-WPRE (196416), AAV-GfaABC1D-MCS--4×6T-WPRE (196417), AAV-CAG-flex-GFP-4×6T (196418), AAV-CAG-flex-lck-smV5-4×6T (196419), AAV-CAG-flex-MCS--4×6T (196420), AAV-CAG-dDIO-lck-smMyc-4×6T (196421), AAV-CAG-dDIO-MCS--4×6T (196422), AAV-CAG-flex-lck-smV5 (196423). The Ribotag input and immunoprecipitated RNAseq datasets generated during the current study are in the process of being submitted to GEO. Due to their large size, imaging datasets are available upon request; source data for imaging datasets are provided with this paper.

**Supplementary Data 1**: RNAseq gene lists: IP samples, FDR < 0.05

**Supplementary Data 2**: GSEA gene sets in viral cohorts, FDR < 0.05

**Supplementary Data 3**: Potential miR target genes, FDR < 0.05

## Acknowledgements

This work was supported by the Dr. Miriam and Sheldon G. Adelson Medical Research Foundation (to S.T.C. and R.K.) and the American Federation of Aging Research (to A.J.G.). We thank the members of S.T.C’s laboratory for helpful discussions, Shutong Hou and Christine Hakobyan for assistance with tissue processing, and Qing Wang for assistance with RNA library preparation. This project involved a wide variety of plasmids and DNA constructs; we thank Gilles Bonvento for kindly sharing the miR124T cassette, and Addgene and their depositors for providing all other plasmids that made this work possible. Schematic diagrams created with Biorender.com.

## Contributions

A.J.G. and S.T.C. designed the experiments. A.J.G. constructed the plasmids, packaged the viruses, performed the experiments, and analyzed the results. R.K. performed transcriptomic data analysis. M.V.S. provided reagents. A.J.G. and S.T.C. wrote the manuscript with contributions from all authors.

## Materials & Correspondence

Correspondence to S. Thomas Carmichael.

---

**Supplementary Figure 1.**
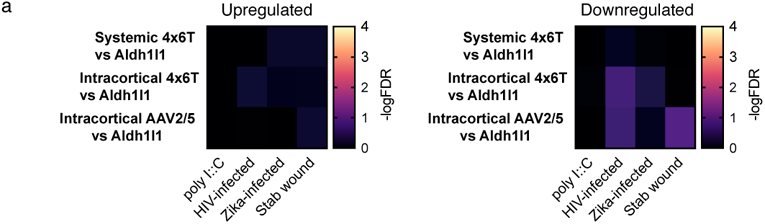
GSEA comparison of astrocytic gene changes induced by AAV transduction vs viral infection or stab wound. **a**, Up- and down-regulated gene sets associated with astrocytic infection by HIV or Zika virus, viral mimetic poly I::C, or stab wound injury were compared with gene changes in AAV-transduced astrocytes (vs Aldh1l1-CreERT2 transcriptomes). None of these gene sets were enriched (FDR < 0.05; −logFDR >1.30103), suggesting that the gene changes induced by AAV transduction are not reminiscent of the gene changes astrocytes undergo in these pathologic conditions.

## References

1. Khakh, B. S. & Sofroniew, M. V. Diversity of astrocyte functions and phenotypes in neural circuits. Nat Neurosci 18, 942–952 (2015).

2. Khakh, B. S. & Deneen, B. The Emerging Nature of Astrocyte Diversity. Annu. Rev.Neurosci. 42, 187–207 (2019).

3. Nectow, A. R. & Nestler, E. J. Viral tools for neuroscience. Nat Rev Neurosci 21, 669–681 (2020).

4. Chan, K. Y. et al. Engineered AAVs for efficient noninvasive gene delivery to the central and peripheral nervous systems. Nat Neurosci 20, 1172–1179 (2017).

5. Griffin, J. M. et al. Astrocyte-selective AAV gene therapy through the endogenous GFAP promoter results in robust transduction in the rat spinal cord following injury. Gene Ther 26, 198–210 (2019).

6. Wang, L.-L. et al. Revisiting astrocyte to neuron conversion with lineage tracing in vivo. Cell 184, 5465–5481.e16 (2021).

7. Le, N., Appel, H., Pannullo, N., Hoang, T. & Blackshaw, S. Ectopic insert-dependent neuronal expression of GFAP promoter-driven AAV constructs in adult mouse retina. Front. Cell Dev. Biol. 10, 914386 (2022).

8. Leib, D., Chen, Y. H., Monteys, A. M. & Davidson, B. L. Limited astrocyte-to-neuron conversion in the mouse brain using NeuroD1 overexpression. Molecular Therapy 30, 982–986 (2022).

9. Chen, W., Zheng, Q., Huang, Q., Ma, S. & Li, M. Repressing PTBP1 fails to convert reactive astrocytes to dopaminergic neurons in a 6-hydroxydopamine mouse model of Parkinson’s disease. eLife 11, e75636 (2022).

10. Guo, T. et al. Downregulating PTBP1 Fails to Convert Astrocytes into Hippocampal Neurons and to Alleviate Symptoms in Alzheimer’s Mouse Models. J. Neurosci. 42, 7309–7317 (2022).

11. Brown, B. D., Venneri, M. A., Zingale, A., Sergi, L. S. & Naldini, L. Endogenous microRNA regulation suppresses transgene expression in hematopoietic lineages and enables stable gene transfer. Nat Med 12, 585–591 (2006).

12. Colin, A. et al. Engineered lentiviral vector targeting astrocytes *In vivo*. Glia 57, 667– 679 (2009).

13. Taschenberger, G., Tereshchenko, J. & Kügler, S. A MicroRNA124 Target Sequence Restores Astrocyte Specificity of gfaABC1D-Driven Transgene Expression in AAV-Mediated Gene Transfer. Molecular Therapy - Nucleic Acids 8, 13–25 (2017).

14. Brenner, M. Structure and Transcriptional Regulation of the GFAP Gene. Brain Pathology 4, 245–257 (1994).

15. Lee, Y., Messing, A., Su, M. & Brenner, M. *GFAP* promoter elements required for region-specific and astrocyte-specific expression. Glia 56, 481–493 (2008).

16. Gao, Y. et al. Develop an efficient and specific AAV-based labeling system for Muller glia in mice. Sci Rep 12, 22410 (2022).

17. Sun, W. et al. SOX9 Is an Astrocyte-Specific Nuclear Marker in the Adult Brain Outside the Neurogenic Regions. J. Neurosci. 37, 4493–4507 (2017).

18. Botterill, J. J. et al. Off-Target Expression of Cre-Dependent Adeno-Associated Viruses in Wild-Type C57BL/6J Mice. eNeuro 8, ENEURO.0363-21.2021 (2021).

19. Madisen, L. et al. A robust and high-throughput Cre reporting and characterization system for the whole mouse brain. Nat Neurosci 13, 133–140 (2010).

20. Hoye, M. L. et al. MicroRNA Profiling Reveals Marker of Motor Neuron Disease in ALS Models. J. Neurosci. 37, 5574–5586 (2017).

21. Jovicic, A. et al. Comprehensive Expression Analyses of Neural Cell-Type-Specific miRNAs Identify New Determinants of the Specification and Maintenance of Neuronal Phenotypes. Journal of Neuroscience 33, 5127–5137 (2013).

22. Fernández-Hernando, C. & Suárez, Y. MicroRNAs in endothelial cell homeostasis and vascular disease: Current Opinion in Hematology 25, 227–236 (2018).

23. Viswanathan, S. et al. High-performance probes for light and electron microscopy. Nat Methods 12, 568–576 (2015).

24. Foo, L. C. & Dougherty, J. D. Aldh1L1 is expressed by postnatal neural stem cells *in vivo*: *In Vivo* Expression of Aldh1L1 by Neural Stem Cells. Glia 61, 1533–1541 (2013).

25. Hochgerner, H., Zeisel, A., Lönnerberg, P. & Linnarsson, S. Conserved properties of dentate gyrus neurogenesis across postnatal development revealed by single-cell RNA sequencing. Nat Neurosci 21, 290–299 (2018).

26. Beyer, F., Lüdje, W., Karpf, J., Saher, G. & Beckervordersandforth, R. Distribution of Aldh1L1-CreERT2 Recombination in Astrocytes Versus Neural Stem Cells in the Neurogenic Niches of the Adult Mouse Brain. Front. Neurosci. 15, 713077 (2021).

27. Zamanian, J. L. et al. Genomic Analysis of Reactive Astrogliosis. Journal of Neuroscience 32, 6391–6410 (2012).

28. Barreto, G. E., Sun, X., Xu, L. & Giffard, R. G. Astrocyte Proliferation Following Stroke in the Mouse Depends on Distance from the Infarct. PLoS ONE 6, e27881 (2011).

29. Ohab, J. J., Fleming, S., Blesch, A. & Carmichael, S. T. A Neurovascular Niche for Neurogenesis after Stroke. Journal of Neuroscience 26, 13007–13016 (2006).

30. Fenno, L. E. et al. Comprehensive Dual- and Triple-Feature Intersectional Single-Vector Delivery of Diverse Functional Payloads to Cells of Behaving Mammals. Neuron 107, 836–853.e11 (2020).

31. Sanz, E. et al. Cell-type-specific isolation of ribosome-associated mRNA from complex tissues. Proc. Natl. Acad. Sci. U.S.A. 106, 13939–13944 (2009).

32. Srinivasan, R. et al. New Transgenic Mouse Lines for Selectively Targeting Astrocytes and Studying Calcium Signals in Astrocyte Processes In Situ and In Vivo. Neuron 92, 1181–1195 (2016).

33. Becher, B., Waisman, A. & Lu, L.-F. Conditional Gene-Targeting in Mice: Problems and Solutions. Immunity 48, 835–836 (2018).

34. Sofroniew, M. V. Molecular dissection of reactive astrogliosis and glial scar formation. Trends in Neurosciences 32, 638–647 (2009).

35. McKenzie, A. T. et al. Brain Cell Type Specific Gene Expression and Co-expression Network Architectures. Sci Rep 8, 8868 (2018).

36. Subramanian, A. et al. Gene set enrichment analysis: A knowledge-based approach for interpreting genome-wide expression profiles. Proc. Natl. Acad. Sci. U.S.A. 102, 15545–15550 (2005).

37. Li, J. et al. Conservation and divergence of vulnerability and responses to stressors between human and mouse astrocytes. Nat Commun 12, 3958 (2021).

38. Shereen, M. A. et al. Zika virus dysregulates the expression of astrocytic genes involved in neurodevelopment. PLoS Negl Trop Dis 15, e0009362 (2021).

39. Edara, V. V., Ghorpade, A. & Borgmann, K. Insights into the Gene Expression Profiles of Active and Restricted Red/Green-HIV ^+^ Human Astrocytes: Implications for Shock or Lock Therapies in the Brain. J Virol 94, e01563–19 (2020).

40. Sirko, S. et al. Astrocyte reactivity after brain injury—: The role of galectins 1 and 3. Glia 63, 2340–2361 (2015).

41. Karagkouni, D. et al. DIANA-TarBase v8: a decade-long collection of experimentally supported miRNA–gene interactions. Nucleic Acids Research 46, D239–D245 (2018).

42. Matsuda, T. & Cepko, C. L. Controlled expression of transgenes introduced by *in vivo* electroporation. Proc. Natl. Acad. Sci. U.S.A. 104, 1027–1032 (2007).

43. Yao, S. et al. RecV recombinase system for in vivo targeted optogenomic modifications of single cells or cell populations. Nat Methods 17, 422–429 (2020).

44. Fenno, L. E. et al. Targeting cells with single vectors using multiple-feature Boolean logic. Nat Methods 11, 763–772 (2014).

45. Cheng, L.-C., Pastrana, E., Tavazoie, M. & Doetsch, F. miR-124 regulates adult neurogenesis in the subventricular zone stem cell niche. Nat Neurosci 12, 399–408 (2009).

46. Chai, H. et al. Neural Circuit-Specialized Astrocytes: Transcriptomic, Proteomic, Morphological, and Functional Evidence. Neuron 95, 531–549.e9 (2017).

47. John Lin, C.-C., et al. Identification of diverse astrocyte populations and their malignant analogs. Nat Neurosci 20, 396–405 (2017).

48. Burda, J. E. et al. Divergent transcriptional regulation of astrocyte reactivity across disorders. Nature 606, 557–564 (2022).

49. Hasel, P., Rose, I. V. L., Sadick, J. S., Kim, R. D. & Liddelow, S. A. Neuroinflammatory astrocyte subtypes in the mouse brain. Nat Neurosci 24, 1475– 1487 (2021).

50. Gibson, D. G. et al. Enzymatic assembly of DNA molecules up to several hundred kilobases. Nat Methods 6, 343–345 (2009).

51. Madisen, L. et al. Transgenic Mice for Intersectional Targeting of Neural Sensors and Effectors with High Specificity and Performance. Neuron 85, 942–958 (2015).

52. Challis, R. C. et al. Systemic AAV vectors for widespread and targeted gene delivery in rodents. Nat Protoc 14, 379–414 (2019).

53. Gombash Lampe, S. E., Kaspar, B. K. & Foust, K. D. Intravenous Injections in Neonatal Mice. JoVE 52037 (2014) doi:10.3791/52037.

54. Li, S. et al. An age-related sprouting transcriptome provides molecular control of axonal sprouting after stroke. Nat Neurosci 13, 1496–1504 (2010).

55. Schindelin, J., et al. Fiji: an open-source platform for biological-image analysis. Nat Methods 9, 676–682 (2012).

56. Sanz, E., Bean, J. C., Carey, D. P., Quintana, A. & McKnight, G. S. RiboTag: Ribosomal Tagging Strategy to Analyze Cell-Type-Specific mRNA Expression In Vivo. Current Protocols in Neuroscience 88, (2019).

57. Vanlandewijck, M. et al. A molecular atlas of cell types and zonation in the brain vasculature. Nature 554, 475–480 (2018).

58. Hulsen, T., de Vlieg, J. & Alkema, W. BioVenn – a web application for the comparison and visualization of biological lists using area-proportional Venn diagrams. BMC Genomics 9, 488 (2008).

59. Liberzon, A. et al. The Molecular Signatures Database Hallmark Gene Set Collection. Cell Systems 1, 417–425 (2015).

